# A Self-Supervised Deep Neural Network for Image Completion Resembles Early Visual Cortex fMRI Activity Patterns for Occluded Scenes

**DOI:** 10.1101/2020.03.24.005132

**Authors:** Michele Svanera, Andrew T. Morgan, Lucy S. Petro, Lars Muckli

## Abstract

The promise of artificial intelligence in understanding biological vision relies on the comparison of computational models with brain data with the goal of capturing functional principles of visual information processing. Convolutional neural networks (CNN) have successfully matched the transformations in hierarchical processing occurring along the brain’s feedforward visual pathway extending into ventral temporal cortex. However, we are still to learn if CNNs can successfully describe feedback processes in early visual cortex. Here, we investigated similarities between human early visual cortex and a CNN with encoder/decoder architecture, trained with self-supervised learning to fill occlusions and reconstruct an unseen image. Using Representational Similarity Analysis (RSA), we compared 3T fMRI data from a non-stimulated patch of early visual cortex in human participants viewing partially occluded images, with the different CNN layer activations from the same images. Results show that our self-supervised image-completion network outperforms a classical object-recognition supervised network (VGG16) in terms of similarity to fMRI data. This provides additional evidence that optimal models of the visual system might come from less feedforward architectures trained with less supervision. We also find that CNN decoder pathway activations are more similar to brain processing compared to encoder activations, suggesting an integration of mid- and low/middle-level features in early visual cortex. Challenging an AI model and the human brain to solve the same task offers a valuable way to compare CNNs with brain data and helps to constrain our understanding of information processing such as neuronal predictive coding.

## Introduction

Investigating the functional dichotomy of forward and backward pathways in the visual system is necessary for describing how the brain performs vision. The conceptual divergence is broadly understood; forward pathways carry unlabelled sensory information into the brain while feedback pathways carry signals from higher visual and non-visual areas back to earlier areas ([Schwiedrzik and Freiwald, 2017, Pennartz et al., 2019]), in the opposite direction to sensory information. Feedback has an important role in the contextual modulation of the feedforward stream, carrying top-down signals such as attention and expectations. ([Van Essen and Anderson, 1995, Gilbert and Li, 2013, Roelfsema and de Lange, 2016, Angelucci et al., 2017, Pennartz et al., 2019, Klink et al., 2017, Morgan et al., 2019, Takahashi et al., 2016]). Neuronal information processing in visual perception can be formalised as statistical inference based on hierarchical internal models, which can theoretically be implemented by schemes such as predictive coding ([Rao and Ballard, 1999, Friston, 2005]). Predictive processing is a compelling framework for describing neural phenomena observed in human primary visual cortex using brain imaging ([Alink et al., 2010, Edwards et al., 2017]). Predictive coding describes a neuronal coding process in which predicted information from top-down streams is explained away from the feedforward stream (e.g., [Rao and Ballard, 1999]). Less narrow implementations of predictive coding recognise the necessity of the feedforward stream to not only compute prediction error but also communicate information that reinforces the internal models which were successful in predicting incoming information ([Shipp, 2016, Rao and Ballard, 1999]). We use the term predictive processing to allow for an even broader implementation of coding principles, including those that provide context (e.g., contour ownership, spatial information from auditory cues) without being limited to predicting a narrow set of precise features (e.g., a specific contour). For example, illusions of motion increase activity on the non-stimulated motion path in primary visual cortex [Erlikhman and Caplovitz, 2017, Chong et al., 2016], and predictable stimuli presented on the motion path cause less activity than surprising stimuli ([Alink et al., 2010, Edwards et al., 2017]). Moreover, visual contours predicted by flanking stimuli increase activity along the illusory contour [Kok et al., 2016, Bergmann et al., 2019]. These data reveal that some neuronal activity in early visual areas is not directly related to retinotopic sensory inputs but rather to the brain’s inference of the world, transmitted in cortical feedback pathways to earlier areas. A comprehensive biologically constrained artificial model of vision should account for this top-down brain processing.

In computational neuroscience, there is a tradition to exploit recent developments in statistical modelling and algorithms to develop more sophisticated models of visual processing ([Simoncelli and Olshausen, 2001]). In this respect, recent developments in artificial intelligence (AI), and in particular deep learning (DL), offered significant contributions in the last decade ([Hassabis et al., 2017]), showing promising results in the attempt to improve our understanding of visual counterstream computations ([Kietzmann et al., 2019, Qiao et al., 2019]). In the DL field, and more broadly in AI, a textbook categorisation in the literature is the distinction between supervised and unsupervised learning; while the first learns statistical representations based on labelled datasets, including tasks such as classification, regression, and segmentation, the latter tries to extract features and inherent patterns from unlabelled data, with tasks such as clustering, anomaly detection, and dimensionality reduction ([Hastie et al., 2009]). While supervised learning usually allows to obtain better performance, being able to make sense of the large amount of unlabelled data often available is a promising and exciting goal for the field. In this respect, the recent trend of self-supervised learning tries to use the data as a supervision strategy ([Jing and Tian, 2020]): the idea is to challenge the model to solve a specific unrelated task in order to learn representations of the data or to automatically label the dataset. Examples include learning to predict some part of the image from other parts, predicting relative locations of two image patches, solving a jigsaw puzzle, and colorizing an image. Originally designed as unsupervised methods, in Generative Adversarial Networks (GAN - [Goodfellow et al., 2014]), a discriminator is trained by evaluating whether the data created by a generator is part of the training data (i.e., is a real image) or not (fake). Despite their great promise of data understanding, unsupervised learning methods are not often used for model comparisons with brain data.

Supervised learning models, under the flagship of Convolutional Neural Networks (CNN), have advanced our understanding of visual information processing in the brain in a number of breakthrough studies ([Hassabis et al., 2017]). Using this approach to model spiking computations, CNN models can predict neural activations in the macaque visual ventral streams at early time points after stimulus presentation, suggesting they may capture important aspects of feedforward visual processing ([Yamins and DiCarlo, 2016]). Natural images have been found to cluster together in similar ways in the internal feature spaces of CNNs as in human Inferior Temporal (IT) cortex ([Khaligh-Razavi and Kriegeskorte, 2014]). [Cichy et al., 2016] described how a CNN captured the stages of human visual processing in time and space from early visual areas towards the dorsal and ventral streams. [Kay et al., 2008] and [Güçlü and van Gerven, 2015] compared two different encoding models to human fMRI data, one based on decomposing visual information into Gabor elements, and the other based on trained feedforward CNNs. Comparisons showed improved results of the supervised CNN with respect to the Gabor based approach in explaining fMRI activity patterns. Using decoding modelling, different studies ([Eickenberg et al., 2017, He et al., 2018]) proposed to derive CNN representations from brain data, or transfer learning CNN discrimination properties for fMRI decoding ([Svanera et al., 2019]). In a broader effort to validate CNNs as models of the visual system, supervised networks on image recognition were used to model differences in retinal and cortical networks showing which architecture constraints help the emergence of similar CNN representations in the brain ([Lindsey et al., 2019]). To better navigate between different object detection artificial networks, Schrimpf et al. ([Schrimpf et al., 2018]) introduce multiple neural and behavioural benchmarks to score any artificial neural network on how similar it is to the brain’s mechanisms for core object recognition. CNNs with recurrent connections better predict visual responses than feedforward models, and increase our ability to capture the cortical dynamics in MEG data and fMRI data ([Kietzmann et al., 2019, Qiao et al., 2019]), opening new opportunities on understanding visual counterstreams by modelling brain architectures and processing principles [Hong et al., 2016, Kar et al., 2019, Tang et al., 2018].

While these achievements are noteworthy in terms of being the best available models of biological vision, the possible impact of unsupervised models remains comparatively under explored. As in AI, unsupervised learning models are promising in terms of data exploitation and understanding. Furthermore, in modelling vision we could potentially narrow the gap created by the biological implausibility of supervised learning objectives and gain progress in understanding the role of cortical feedback processing. In addition, the setting (i.e., the task) in which the two models, biological and artificial, are compared could provide a better testbed and bring potential benefits, such as giving insights on the learning dynamics or identifying differences with respect to other tasks. We begin in this direction by comparing brain data capturing contextual feedback in vision (in amodal filling-in) to an artificial model trained to fill-in missing visual information. The human brain imaging data was recorded from primary visual cortex (V1) whilst subjects viewed partially occluded natural scene images ([Smith and Muckli, 2010, Muckli et al., 2015, Revina et al., 2018, Morgan et al., 2019]). By partially masking the visual stimulus and recording from the corresponding retinotopic region of V1, this paradigm allows us to study feedback information in the absence of stimulus-specific feedforward signals. We use this data as a testbed for investigating a vision model that includes feedback components. We selected a visual occlusion paradigm because we sought comparable functionality between the brain data and computer vision algorithms performing the same task. The predictive coding framework suggests that the brain is trying to reconstruct the image under the occlusion, using information from cortical feedback to infer the content of missing image sections. We showed recently that the brain’s filling of the missing image section can be modelled as a behavioural line drawing [Morgan et al., 2019]. This operation of reconstructing an unseen image is conceptually similar to the task of inpainting (also known as image completion) in computer vision. In this task, an artificial model predicts the missing part of the image (unseen or damaged, [Pathak et al., 2016]) relying on image statistics learnt during training. This learning can be obtained using different techniques: with classical image processing techniques, as the application of statistically similar patches ([He and Sun, 2014]), or with CNN approaches ([Yu et al., 2018, Lempitsky et al., 2018, Isola et al., 2017]). To solve the task of image reconstruction, we used a CNN with encoder/decoder architecture, trained in a self-supervised fashion as in [Isola et al., 2017], to perform inpainting on the lower-right quadrant i.e., reconstructing the unseen (occluded) portion of the image (Figure 1 - a). We compared the encoder/decoder network trained to solve inpainting with brain data collected in a previous 3-Tesla functional magnetic resonance imaging (fMRI) experiment during the viewing of images with the lower-right quadrant occluded (Figure 1 - b), see [Morgan et al., 2019]). We investigated similarities between the network and brain data using Representational Similarity Analyses (RSA, [Kriegeskorte et al., 2008, Nili et al., 2014]).

**Figure 1:**
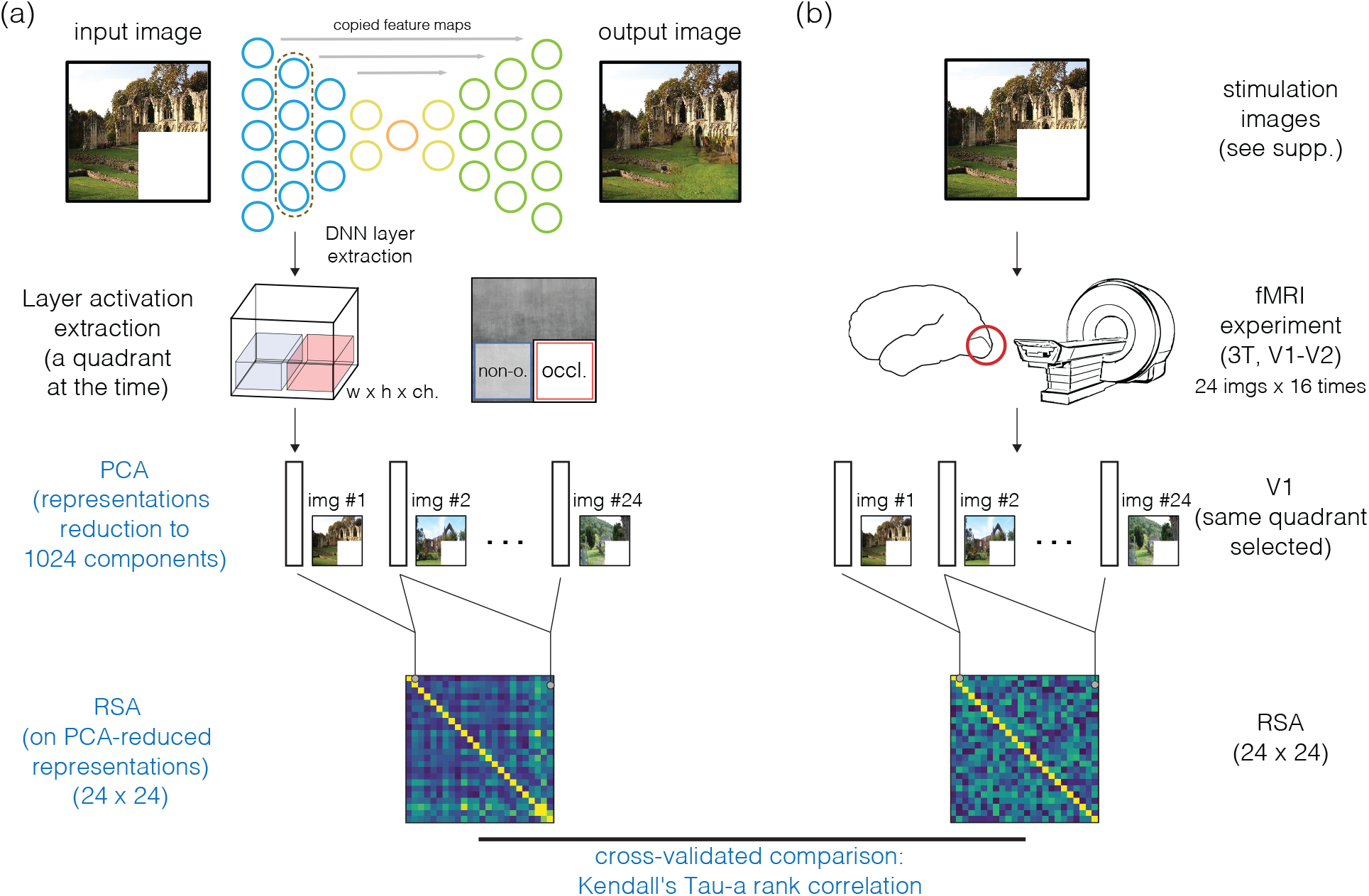
The analysis framework is composed of two parts: the encoder/decoder (artificial) neural network and brain imaging data collection. (a) The image passed through the network, we extracted activations for one layer, we selected a quadrant at the time, and we applied PCA transformation to reduce the dimension to 1024 components; we then obtained one 1024-d vector per layer (15), per quadrant (2), and per image (24). We used these vectors to compute Representational Dissimilarity Matrices (RDMs). (b) fMRI data were collected from participants whilst viewing the same images that were fed into the network (in testing) and RDMs computed. We compared RDMs of the network and brain data (cross-validated - see [Walther et al., 2016]), for every CNN layer (15 layers analysed), human visual area (V1 and V2), and image space quadrant (occluded and non-occluded quadrants).

## Material and methods

We compared representations of two neural network models to brain imaging data acquired during an fMRI experiment. We first describe the fMRI experiment (Section *fMRI experiment*), before specifying the two CNN models: the well-known supervised (feedforward) network VGG16 and our encoder/decoder model trained to fill the missing quadrant of the image (Section *Artificial neural network models*). Lastly, we explain how we compared the brain data to the CNNs usingRepresentational Similarity Analysis (RSA) (Section *Data analysis: Representational similarity analysis (RSA)*). The experimental framework is shown in Figure 1.

### fMRI experiment

Our fMRI dataset was collected previously and published in [Morgan et al., 2019]. Eighteen healthy volunteers with normal or corrected-to-normal vision participated in this study. Twenty-four real-world scenes from six categories (beaches, buildings, forests, highways, industry, and mountains), from the dataset in ([Walther et al., 2009]) were shown to participants ([Morgan et al., 2019]; see supplementary for stimulation images). Images were displayed in grayscale on a rear-projection screen using a projector system. Stimuli spanned 19.5° × 14.7° of visual angle and were presented with the lower-right quadrant occluded by a white box (occluded region spanned ∼ 9° × 7°). A centralised fixation checkerboard marked the centre of the scene images. Over the course of the experiment, each image was presented 16 times. Refer to [Morgan et al., 2019] for further details^2^.

### Data acquisition

We collected the fMRI data previously at the Centre for Cognitive Neuroimaging, at the University of Glasgow. Participants gave written informed consent to participate, in accordance with the institutional guidelines of the local ethics committee of the College of Science & Engineering at the University of Glasgow (CSE01127). We used EPI sequences to acquire partial brain volumes aligned to maximise coverage of the visual pathway (18 slices; voxel size: 3mm, isotropic; 0.3mm interslice gap; TR = 1000ms; TE = 30ms; matrix size = 70 × 64; FOV = 210 × 192mm). We used retinotopic mapping data ([Sereno et al., 1995, Wandell et al., 2007]) to locate V1 and V2 (polar angle and eccentricity mapping). Additionally, we used three flashing checkerboard locations to map cortical subregions of V1 and V2 corresponding to occluded and non-occluded portions of the visual field. The first checkerboard was located in the bottom-right visual quadrant (‘Target mapping’ condition), the second was located only at the border (‘Surround mapping’ condition; extending 2° visual angle into the occluded quadrant) and the third covered the remaining three quadrants (‘Non-Occluded mapping’ condition). We declared cortical subregions that responded statistically more to the contrast of ‘Target’ vs. ‘Surround’ conditions as occluded i.e., not stimulated with scene information. In our non-occluded condition, the visual field was presented with scene information, activating the corresponding retinotopic region of V1 and V2.

### Data pre-processing

Functional data passed through different pre-processing steps; slice time, 3D motion correction, and temporal filtering (high-pass), before being normalised to Talairach space. We used data from retinotopic mapping runs to define early visual areas V1 and V2 using linear cross-correlation of eight polar angle conditions. To define the regions of interest (ROI) corresponding to the non-occluded and occluded quadrants, lower left and right image sections respectively, we computed population receptive fields (pRF, [Dumoulin and Wandell, 2008]) and excluded voxels whose response profiles were not fully contained (within 2*σ* of their pRF centre) by the respective visual ROI. The patterns of brain activities we obtained, which consisted of data from V1 and V2 for each of the two image quadrants analysed, were not pre-processed with any dimensionality reduction technique (contrary to CNN activations that went through a PCA analysis, see below). We discarded the upper quadrants due to the visual content of stimulation images (shown in supplementary). Most of the images included depict landscapes, and the upper parts of the images therefore included large portions of sky (i.e., uniform values).

### Artificial neural network models

#### VGG16: a supervised image classification network

To compare brain data with a supervised network, we used the VGG16 network ([Simonyan and Zisserman, 2014]), a well-known model used for comparing visual pathway activity with CNNs ([Cichy et al., 2016, Güçlü and van Gerven, 2015]). The network is a supervised model ([Simonyan and Zisserman, 2014]) pre-trained to solve image classification - the task to attribute an object class label to an image - on the ImageNet database with 1000 classes[Russakovsky et al., 2015]). The architecture is strictly hierarchical - image-to-labels - with 16 convolutional layers, and it uses repetitive and simple base structure made by 3 × 3 convolutional layers, for a total of ∼ 138M parameters. In our analyses, we used only convolutional layers before pooling, leading to a total of 5 layers analysed (conditions: `conv1_2`, `conv2_2`, `conv3_3`, `conv4_3`, `conv5_3`).

#### Encoder/decoder model: a self-supervised image completion network

Our self-supervised image-to-image model is trained to reconstruct the occluded image section (always the lower-right quadrant); it is a fully-convolutional neural network, with a encoder/decoder architecture and with skip connections (known as U-Net and described in [Ronneberger et al., 2015]). Details on layer activation dimensions are shown in Figure 2. Every layer implements the same components: a 2-D convolution with a 3 × 3 filter and stride= 2 (that downsamples the activations, no pooling used), batch normalisation, and a rectified linear unit (ReLu) function for encoder and a leaky ReLu^3^ for decoder. We added padding to keep the activation dimensions constant after convolution and adopted dropout during the training to reduce overfitting.

**Figure 2:**
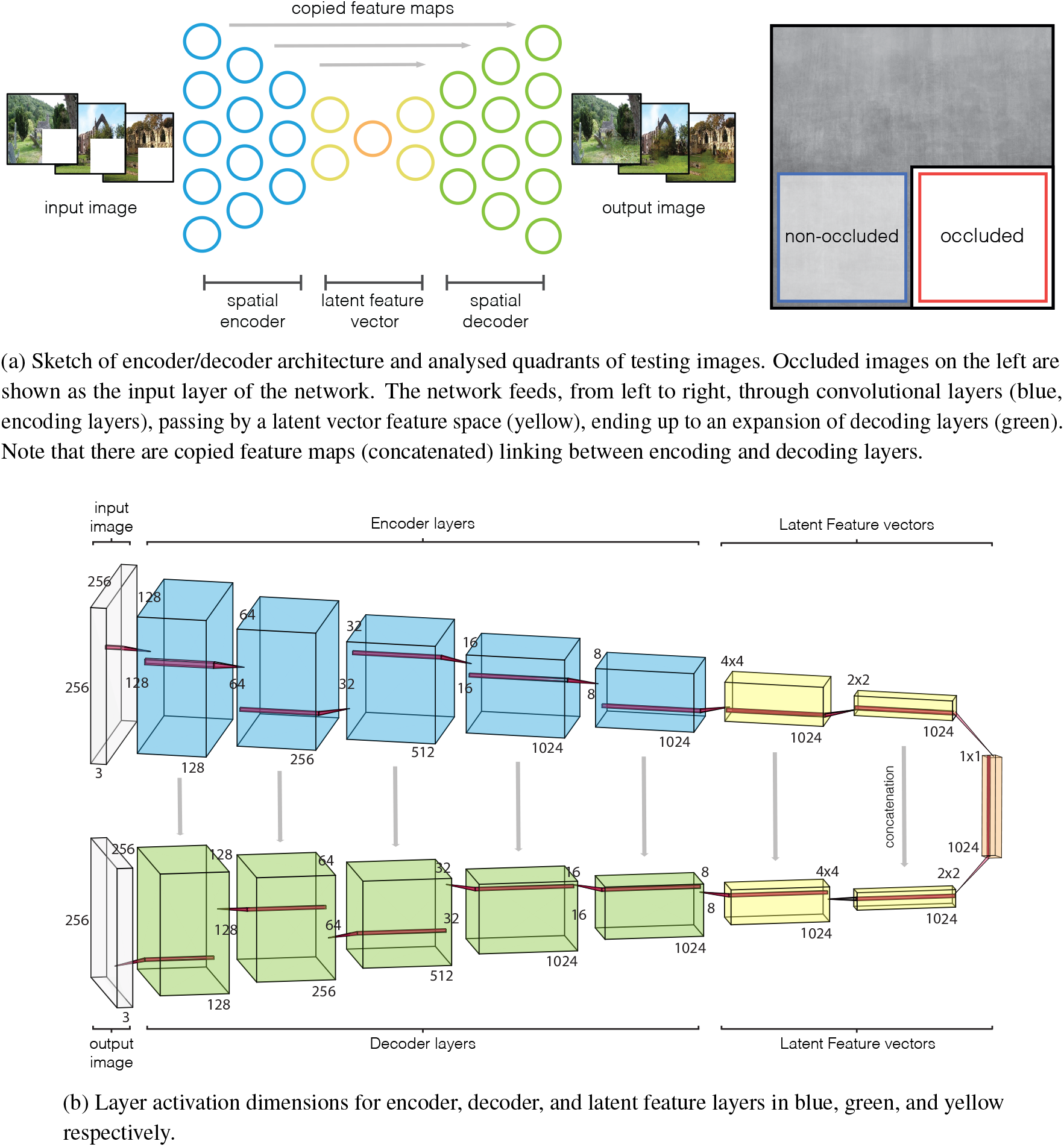
Model architecture with layer dimensions. Every layer implements: convolution, batch normalisation, and activation function (leaky relu for encoder and relu for decoder).

We trained the network similarly to what has been proposed in [Isola et al., 2017], in which not only the mapping from input image to output image (i.e., network filters) is learned, but also the best loss function to evaluate (or train) this mapping. The selection of the learning strategy, and in particular the loss, is crucial in unsupervised learning, where labels or categories are not available. Early unsupervised models adopted custom loss functions hand-crafted by the experimenter; for example, autoencoders (AE) use the mean squared error (MSE) between the original image and the generated to compute the loss and the relative gradient. Variational autoencoders (VAE) ([Kingma and Welling, 2013]), a combination of AE with statistical inference, add a term to the MSE loss measuring how closely the latent variables match specific distributions (ex. unit gaussian). However, MSE produces blurry images and VAEs fail to generate good-looking images since they are not able to parametrise precisely the complex distributions of images. A successful approach to overcome the hand-crafted selection of a loss function is to let the network learn it: this can be accomplished using conditional generative adversarial neural networks ([Goodfellow et al., 2014]). The training was carried out using images from the SUN database ([Xiao et al., 2010], the same database where the testing images came from), with occluded images as input to the network and original images (without occlusion) as output. We discarded some images from the database because they were too low resolution (less than 200 × 200 pixels), leading to a total of ∼ 107000 images for the training^4^ . We trained the model and its 228M parameters with epochs= 5 and batch_size= 5 and the training process lasted for ∼ 49 hours on a GeForce 1080Ti. All the code is implemented in tensorflow. An overview of the training and testing procedures is shown in Figure 3 of Supplementary material.

**Figure 3:**
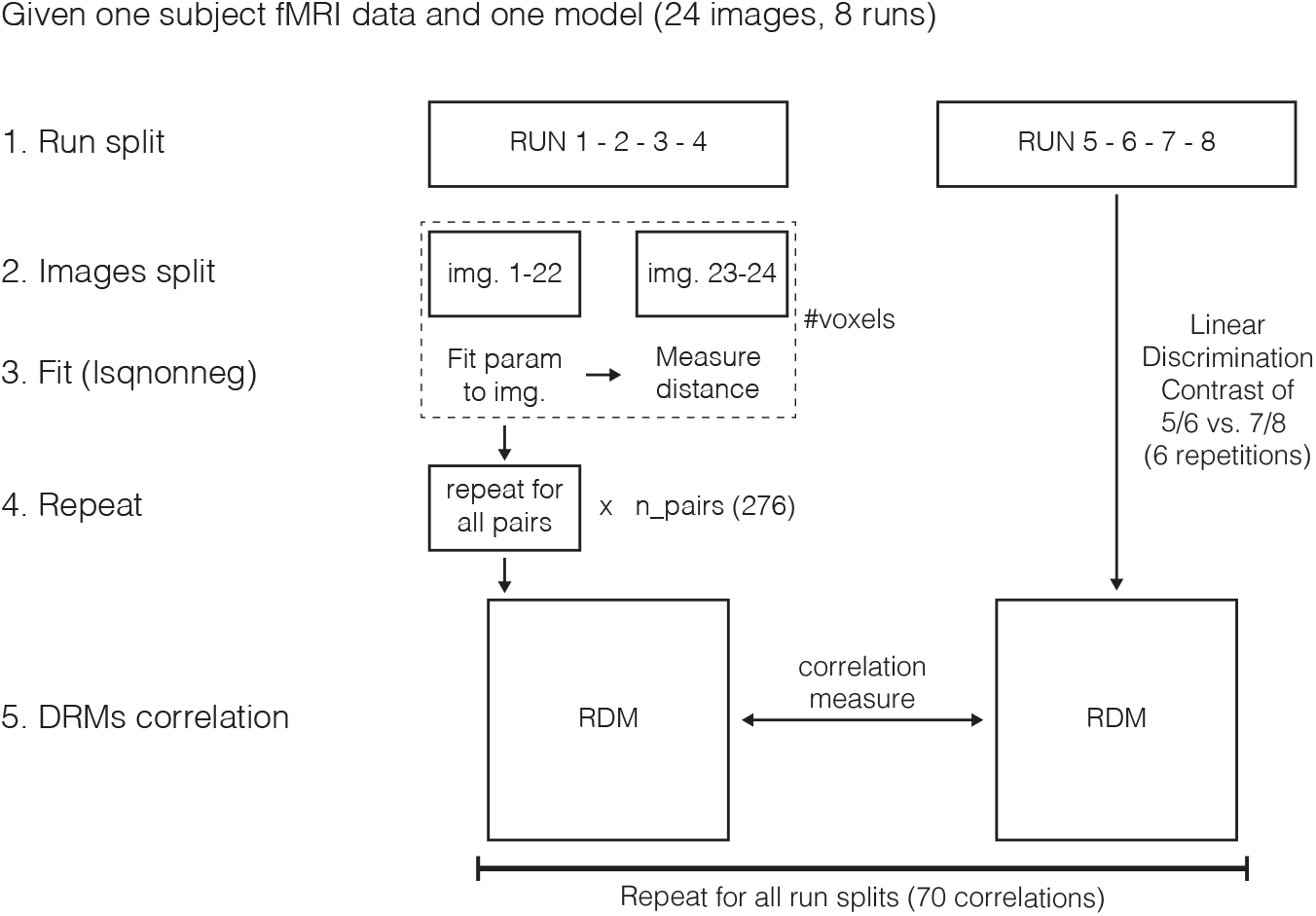
A schematic overview of the RSA computation.

We extracted layer activations from both encoder and decoder streams, as shown in Figure 1 for two quadrants of the image: lower-left (non-occluded), and lower-right (occluded). The contractions in the architecture produce an increase of the (CNN) receptive fields as the layers increase, causing overlaps between quadrants. To avoid confounding results, we selected only an area within the quadrants, removing borders between them. Calculating the receptive field of the network (see supplementary for receptive field sizes), we noted how after the fifth encoder layer, the receptive field size (i.e., the region of the input image which contribute to filter activations) spanned the entire image, and layers representations no longer contained spatiotopic activity maps (this was verified displaying activation maps). It was then not possible to distinguish quadrants by spatial location, so we decomposed the analysis into three conditions: the *spatial encoder*, in which activations could be split into quadrants; *latent feature vectors*, which included activations about the context only and without the spatial information; and *spatial decoder*, with again the concept of spatial separation in reconstruction.

### Dimensionality reduction of layer activations: PCA

As described next (Section *Data analysis: Representational similarity analysis (RSA)*), we performed Representation Similarity Analysis (RSA) through predictions of similarities between CNN and early visual cortex representations. Since layer activations from the CNN have large dimensions (see sizes in Figure 2-(b)), we applied a dimension reduction technique to obtain stable and feasible RSA predictions in terms of variance explained (avoiding overfitting) and computation time. We therefore applied Principal Component Analysis (PCA) to every layer activation of the set.

In order to obtain comparable comparisons in the RSA analysis, we reduced every layer activation to the same dimension, for every layer analysed, as done in [Cichy et al., 2016]. The dimensionality reduction is learnt randomly selecting 10000 images from the training set, extracting the corresponding activations, learning the transformation through an incremental PCA ([Ross et al., 2008, Pedregosa et al., 2011]), and eventually applying the transformation on activations extracted from the testing set. We tested different values for the number of principal components (within 2^*n*^, with *n* = 3, 4, … , 10, i.e., 8, 16, … , 1024 components), as reported in Section *Dimensionality reduction: PCA*.

### Data analysis: Representational similarity analysis (RSA)

We computed Representational Dissimilarity Matrices (RDMs) from fMRI data and PCA-reduced CNN activations using the Linear Discriminant Contrast method ([Walther et al., 2016]). For each scene comparison, we fit models to the RDM of the 22 other scenes in the first set (e.g., runs 1*/*2 vs. 3*/*4) using non-negative least-squares ([Khaligh-Razavi and Kriegeskorte, 2014]). We repeated this for all scene comparisons, producing a predicted RDM based on model parameters, which was then compared to an RDM produced from the other half of the dataset (e.g., runs 5*/*6 vs. 7*/*8) using Kendall’s Tau-a rank correlation ([Nili et al., 2014]). We repeated this procedure for all 70 possible split-quarter combinations, and averaged values over splits to produce one correlation value per subject per ROI and model combinations ([Morgan et al., 2019]). A brief outline of the computation is shown in Figure 3.

## Results

Firstly, we show the visual results of the network output and a description of what layer activations represent (Section *DNN model training*). Then we show similarity performance which drove the selection of the number of PCA components (Section *Dimensionality reduction: PCA*). Lastly, we compare VGG and our encoder/decoder in terms of RSA and encoder/decoder layers (Section *Comparison between VGG16 and encoder/decoder*).

### DNN model training

A graphical result to demonstrate the quality of the output of the network is shown in Figure 4.

**Figure 4:**
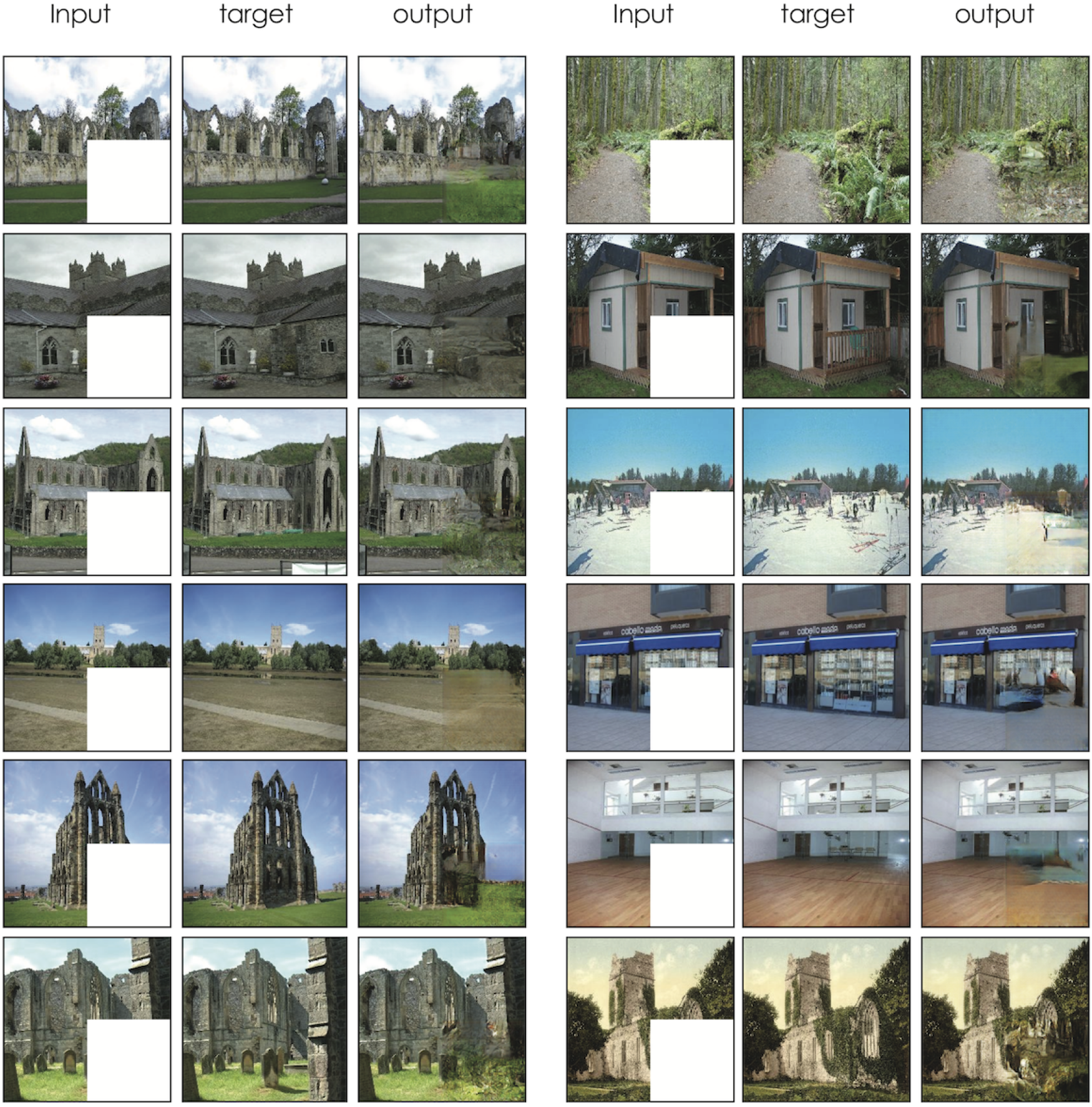
Model results with the input to the network (occluded images), the target used to train the network (i.e., the original image), and the output of the network. Images used in this figure were not part of the training set.

In order to create a model that might capture fMRI brain activation to occluded regions of images, we trained a network to reconstruct grayscale images with occlusion. Images used in the experiment were not part of the training set.

To better understand the processing performed by the network, we show a selection of stimuli eliciting maximal activations for every layer of the network encoder in Figure 5. Patches were obtained as follows: given activations for a specific channel^5^ from 10000 images of the training set, we found the top five activations (maximum responses for that filter). Starting from the location of the maximum, we found the corresponding patch (the receptive field) in the image space that caused that activation. In the classical hierarchical structure of different feedforward networks, such as VGG16 ([Simonyan and Zisserman, 2014]), it is possible to find features from low to high level of complexity. In this network, instead, features appear to be more dedicated to the detection of edges even in middle and upper layers. Only in layer `encoder_6` to `encoder_8` do features start to become more complex, incorporating larger areas of the scene.

**Figure 5:**
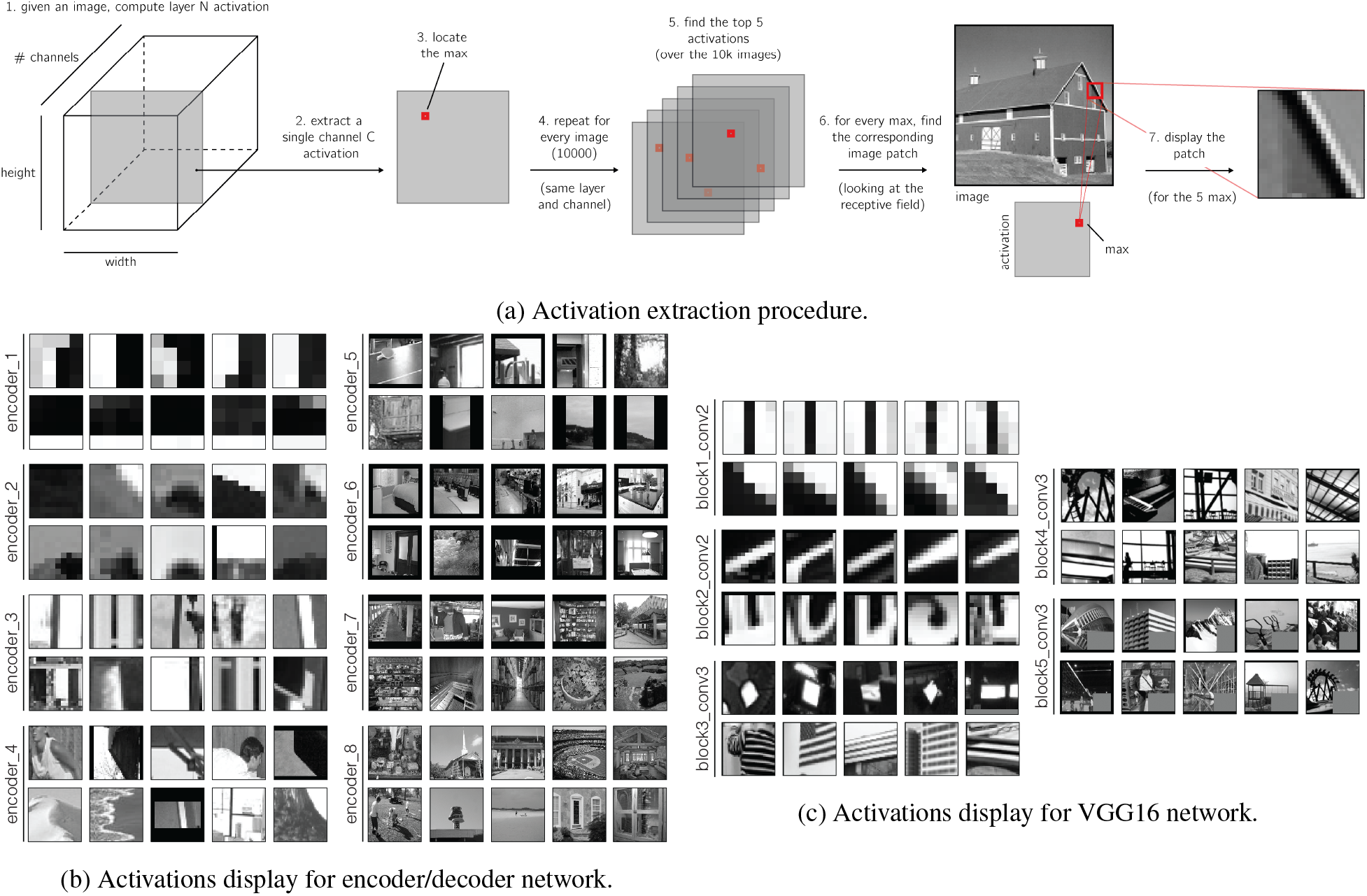
For every layer of a network, two channels are displayed (a row each). The five images correspond to the top five activations for a specific channel, highlighting common visual structure. These images are randomly selected from a bigger pool of analysed layers - the remaining are displayed in the Supplementary Material.

The functional characterisation of these blocks can be challenging, but a possible interpretation follows. The model architecture is composed of two branches: encoder and decoder, also known as contraction and expansion paths. The encoder takes in an original image and produces a compressed vector, or latent variable vector; the decoder takes this vector and tries to reproduce the original image. The encoder branch increases, layer by layer, the representation of “what”, i.e., the feature complexity, and reduces the “where”, i.e., the localisation of a specific feature in the image space. In terms of early visual processing, this is equivalent to compressing retinotopic features into high-level representations.

In contrast, the expansion path creates a high-resolution image on the output through a sequence of up-convolutions (i.e., transposed convolution) and concatenation with high-resolution features from the contracting path, i.e. reconstructs retinotopic features from high-level representations. We chose a contraction (similar to what happens with CNNs for object recognition [Krizhevsky et al., 2012]) and an expansion path, in order to accomplish increased computational efficiency, rather than learning a full-size convolutional net, forcing the network to learn features at different scales ([Noh et al., 2015]).

### Dimensionality reduction: PCA

Considering the type of RSA conducted (Section *Data analysis: Representational similarity analysis (RSA)*), in order to avoid overfitting and reduce the amount of regularisation needed, we reduced the layer activation dimensions to a lower dimension. Therefore, we applied different numbers of principal components (8, 16, … , 1024), and tested the performance achieved (in terms of similarity, Figure 6). The results describe the mean and the standard deviation of similarity across every layer analysed - between V1 and CNN layer RDMs - for every number of components, in terms of Kendall’s tau-a. Results are averaged across all subjects. The two conditions are non-occluded and occluded: the difference is the relative portion of the cortex analysed through the selection of the correspondent receptive fields (pRF analysis). In the non-occluded condition, we analysed the corresponding region of retinotopic cortex that received feedforward stimulation from the image; in the occluded, the region of cortex that processed only the occluder.

**Figure 6:**
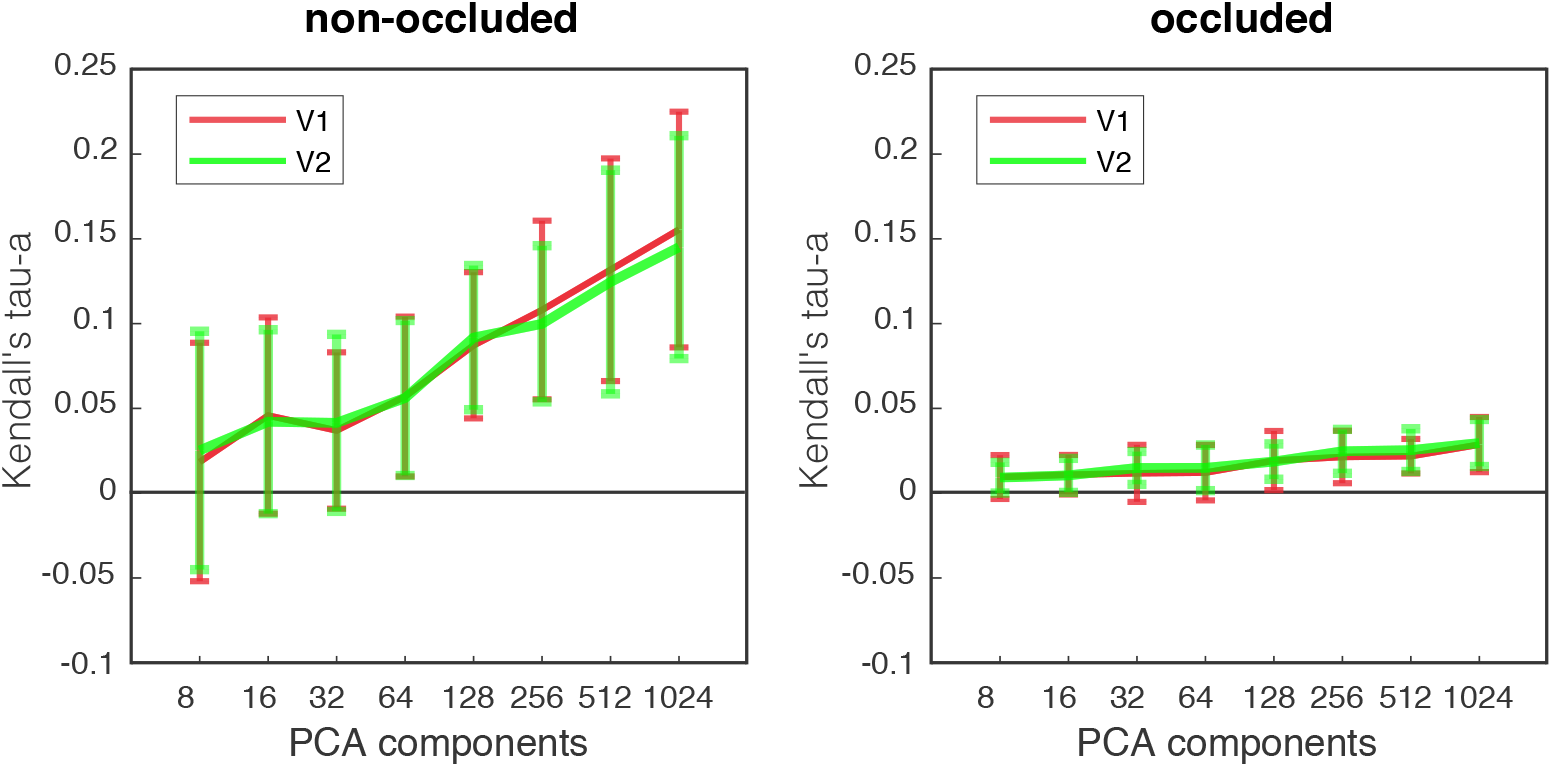
Similarity results (between V1 and CNN layer RDMs) for the non-occluded and occluded quadrants in terms of Kendall’s tau-a. For every component in the range 2^*n*^, with *n* = 3, 4, …, 10, we report mean similarity and standard deviation averaged across layers and subjects.

As expected, with increasing the number of components, correlations become higher and more stable for both V1 and V2. Based on this finding, we kept the number of components equal to 1024. This allowed us to explain the most possible variance in CNN layer activations. Note that 1024 is the maximum reachable number of components, as it is equal to dimensionality of the smallest layer (see Figure 2-b). All results are reported using this number of components; results for VGG are here omitted for brevity, but the same logic is applied.

### Comparison between VGG16 and encoder/decoder

We assessed the ability of the two models to describe brain data using RSA. The first was VGG16 - a supervised network designed for image classification. The second was our encoder/decoder scheme - a model trained in a self-supervised fashion to fill in an occluded portion of images. In Figure 7 we show VGG16 and encoder/decoder response similarities with V1 and V2 in terms of Kendall’s Tau-a rank correlation.

**Figure 7:**
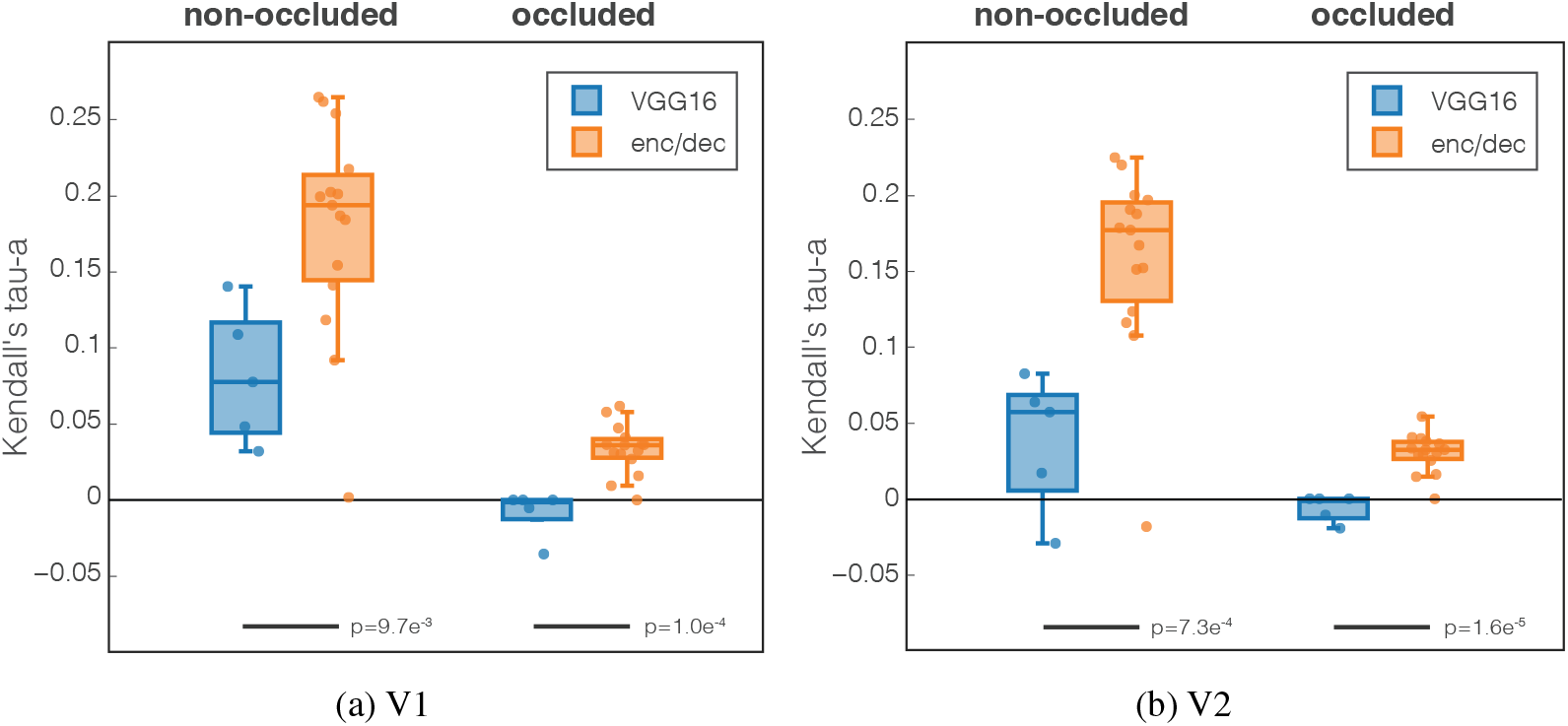
Kendall’s Tau-a rank correlation between VGG16 (blue) and our model (orange) with visual areas (a) V1 and (b) V2. Every dot is a CNN layer (5 conv for VGG16 and 15 for encoder/decoder); results are averaged across subjects. We performed a t-test to determine statistical significance of the difference between the models; results are reported below the bars.

Focusing on the occluded quadrant, we observed a significant difference between VGG16 and our model [*p* <= 0.0097] (differences within our model between encoder and decoder are reported below). This result shows how our model, trained in a self-supervised way to perform inpainting, has representations more similar to the brain data compared to

VGG16’s, a trained supervised model for object recognition. In the non-occluded area, a cortical region combining feedforward and feedback processing, we notice a remarkable increase of similarity between the brain data and our model. We therefore have greater similarity to brain data with our network representations rather than VGG16, for both occluded and non-occluded areas.

### Encoder/decoder layers detail

Focusing on our model, we report results for every layer in Figure 8, analysing differences between encoder (or contraction path) and decoder (or expansion path) parts. We decompose results in the three conditions, *spatial encoder* (blue), *latent feature vectors* (yellow), and *spatial decoder* (green), as described in Section *Encoder/decoder model: a self-supervised image completion network*.

**Figure 8:**
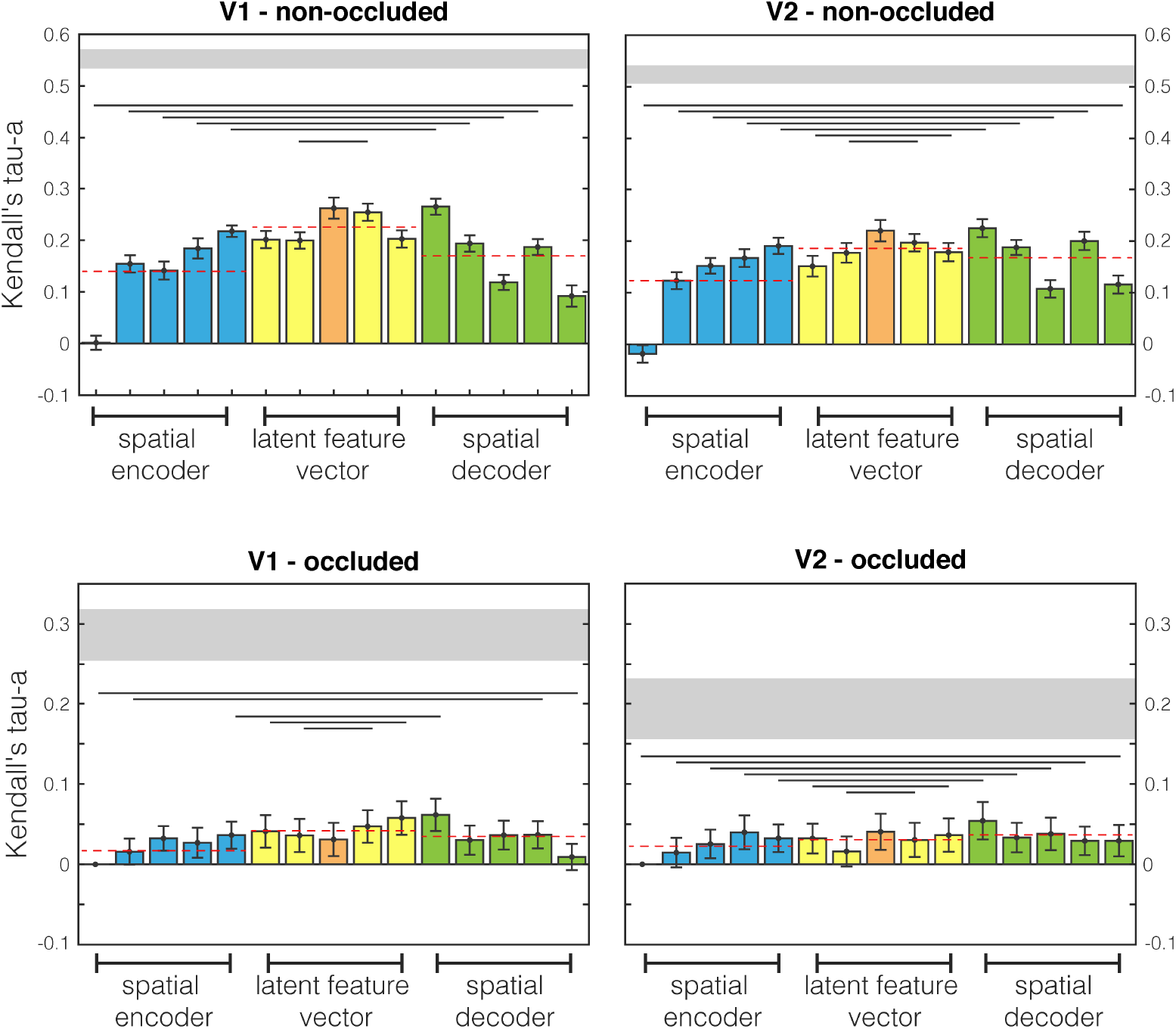
Comparison between brain and encoder/decoder network RDMs. First and second rows are different human visual areas, V1 and V2 respectively. First and second columns relate to non-occluded and occluded quadrants. Results are highlighted with different colours for spatial encoder, latent features, and spatial decoder conditions. Dashed red lines are mean values for the three conditions. Lines below bars show when the encoder and decoder populations difference is significant (Wilcoxon signed rank test).

Four plots are shown: the first row displays V1 results and the second row displays V2; the two columns correspond to non-occluded and occluded quadrants. Every bar is an analysed network layer and colours highlight the three different conditions: encoder (blue), latent feature vector (yellow, with the middle layer in orange), and decoder (green). The bar indicates the mean value across subjects, and the error bar the standard error (standard deviation divided by the square root of the number of subjects - i.e., 18).

Grouping layers together we can highlight large differences between conditions. As depicted by dashed lines, decoder layers (which integrate information from the latent vector space-middle level features- and the correspondent encoders-lower level features-) have greater similarity to brain data with respect to encoders. Surprisingly, the latent feature vector is even higher than decoder, showing the highest similarity with brain data. Analysing layer by layer, we notice that progressive encoding layers (blue bars) become increasingly better descriptions of brain activity patterns. Once the encoder is devoid of retinotopic organisation due to the absence of spatial information, there are the latent feature vectors (yellow bars), which only have “context”, or higher-level features. With these layers, we have an immediate drop in performance with the first layer after the retinotopic layers, which often follows an increase of similarity. In particular, it seems that in the occlusion, for V1, there is a decrease and increase of performance. The orange bar represents the inner layer activation (higher-level) that in some cases represents the higher activation among the context layer. This is until the decoder layers with spatial information (green bars), in which the network reobtains retinotopic organisation. Here, after an initial increase, there is a stabilisation of similarity with brain activity patterns.

Interestingly, the first decoder layer is always the most similar to brain activity patterns. This is the first retinotopically organised layer after integrating the “context” information from latent feature layers. This surprising result opens a series of possibilities for future modelling of brain data from early visual cortex using deep neural networks, such as using a multi-scale approach or adding more complexity than simple edge detection to model V1 processing.

### Receptive field analysis

We investigated whether there was a relationship between the size of V1-V2 receptive fields and encoder/decoder network receptive fields. In Figure Figure 9, we show the receptive field analysis for (a) the CNN encoder layers and (b) the fMRI 3T data. The term receptive field (RF) has here two different meanings, based on the data on which it refers to. On fMRI data, it specifies the size of the visual field that each voxel captures; this is why voxels processing the fovea region have smaller RF size than the peripheral ones. Instead, in CNNs, every unit in a convolutional layer only depends - and processes - a specific region of the input image, called the receptive field. Note that every network layer processes the output of the previous layer, not the original image. In the encoder branch, since “arriving” skip connections are not present (only “departing”), every layer elaborates already processed information because this branch is a cascade of consecutive layers.

**Figure 9:**
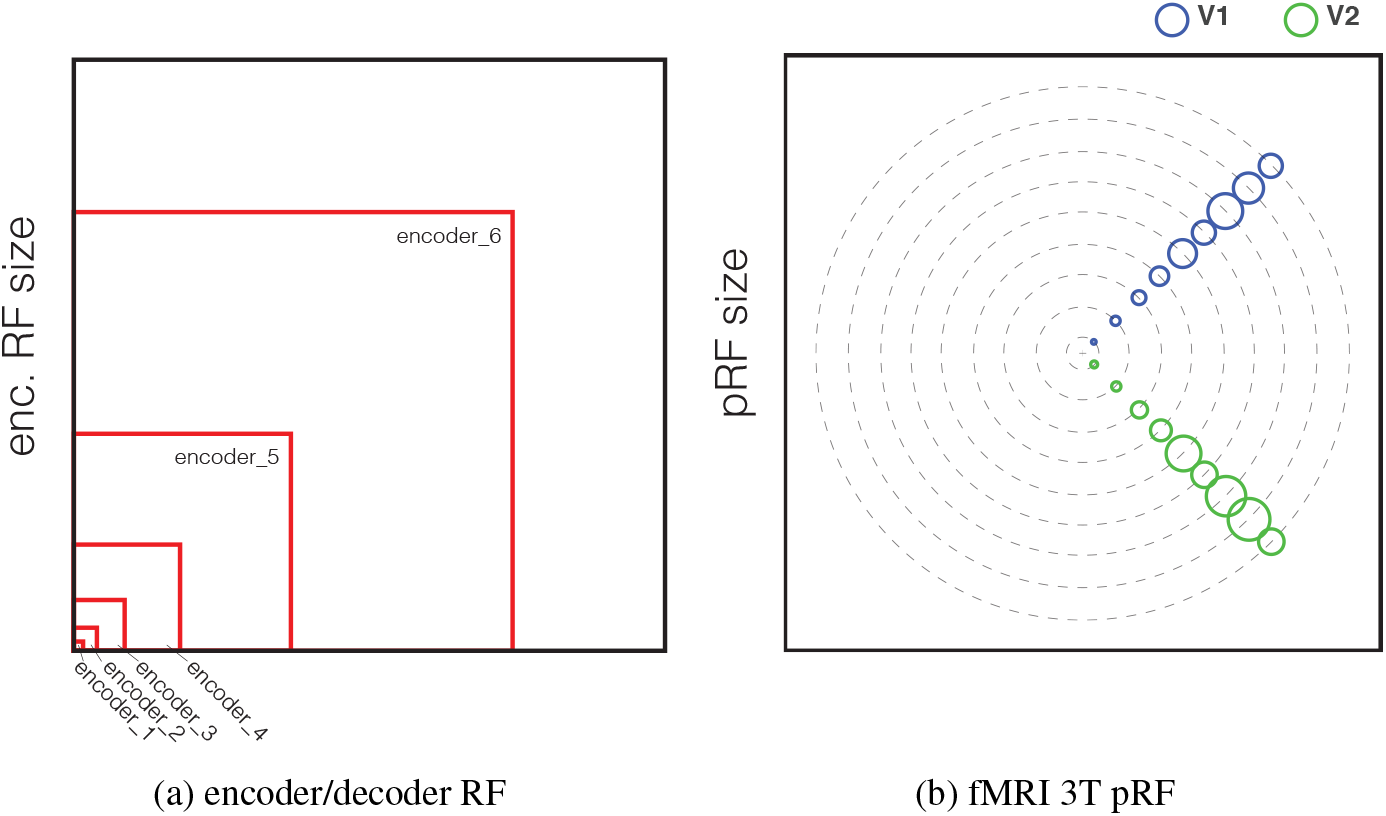
Receptive field analysis. (a) - Receptive field size for the network, encoder layers only, and for (b) the fMRI 3T data. See supplementary for the pRF analyses.

In the Figure 9-(b), we can observe how the RF size of the fMRI data is increasing from the fovea to the periphery (we instructed participants in a central fixation task during the fMRI experiment), for both V1 and V2 (range = {3, 16} pixels for V1 (blue) and {3, 21} for V2 (green)). These data show high similarity with the RF sizes of the first three network layers (`encoder_1` to `encoder_3`, respectively 4 to 22 pixels, see Supplementary for a full list). This may appear in opposition with the results shown in Figure 8, where the similarity increases going from `encoder_1` to `encoder_6`. This means that the increase of similarity further in the network is not due to the matching of the receptive field sizes, but to something else. We can read this result as an evidence of the integration of mid- and low/middle-level features in early visual cortex, since in the decoder layers there is an integration of two different stream of processing: bottleneck and skip connections information.

## Discussion

We investigated the similarity in representations between an artificial neural network trained to fill occlusions and early visual cortical fMRI activity acquired from humans viewing occluded images. Specifically, we wanted to investigate whether an artificial neural network with encoder/decoder architecture would better approximate brain data acquired during a task involving cortical feedback signals, than a purely feedforward network. We have shown two main findings. First, training an artificial network to solve the same task as humans revealed increased similarity to brain data compared to a generic supervised network for classification. Furthermore, the CNN decoder pathway was more similar to brain processing than the encoder pathway. In fact, there was an increase in similarity going further along the network, in line with the integration of multiple feature levels in early visual cortex.

### Comparison between VGG16 and encoder/decoder network

As already speculated ([Hassabis et al., 2017]), a possible common ground for the AI and neuroscience communities is to challenge AI to replicate brain processing while performing common tasks. Having an AI solution could thus lead to better models of the brain for those tasks. With this idea in mind, we trained a network to solve inpainting and compared it to human brain activity performing the same operation. We first compared a supervised network for image classification (VGG16) and an unsupervised network trained on the same task (our model) in terms of brain similarity. In both occluded and non-occluded image locations, we obtained a higher similarity between our model activations and fMRI data; an expected result, considering the features learnt by the CNN. We show in Figure 7 how our model outperforms VGG16, meaning that our model represents images more similarly to the brain in both non-occluded and occluded quadrants.

### Encoder/decoder layers detail

What can our model tell us about brain processing? Our model attempted to reconstruct the occluded image section (lower-right quadrant) using contextual information from the image surround (i.e., the remaining three quadrants) through several layers of processing. The model is composed of two branches, encoder and decoder, which analyses images and reconstructs the missing part in the decoder. The network mainly learns low-level features, including edge detection and texture pattern recognition, at different resolutions (receptive fields), since these are the features considered useful to solve the task (i.e., reconstructing the image).

Going further with the analysis of our network, we tested whether the encoder (i.e., the contraction path) or the decoder (i.e., the expansion path) was more similar to the brain’s representations. We found that the branch of the network more similar to the early visual cortex was not the portion compressing retinotopic features into higher-level representations (i.e., the forward encoder pathway), but instead the portion reconstructing retinotopic features from higher-level representations (i.e., the decoder pathway). Predictive coding theories ([Friston, 2008]) suggest that the brain is trying to reconstruct the image under occlusion, using information in cortical feedback to probabilistically infer the content of missing image sections. Partial or complete occlusion, of objects for example, is commonplace in our visual environments. Predictive coding offers a neuronal mechanism by which the brain can minimise surprise by facilitating the negotiation between top-down predictions and feedforward input once the occlusion is removed. A small warning signal in a cluttered environment can become better detected if predicted information is filtered out. Consistent with such a neuronal framework, our data confirms some level of filling-in with information related to inpainting in computer vision.

### Future directions

In future, it will be important to continue to improve network descriptions of the human visual system by employing more descriptive and specialised networks. Relatedly, we recently showed how line drawings provide a good description of internal model structure representing scene-specific features in human early visual cortex [Morgan et al., 2019]. Being able to build a network able to predict sketches from an image would therefore provide additional value when comparing CNNs to brain data. In computer vision, hand sketches have been extensively studied, for example in sketch recognition, generation, and sketch-based image retrieval [Riaz Muhammad et al., 2018], and reveal that computers can classify line drawings in addition to digital images.

In terms of improving the biological plausibility of neural networks, recent work has also shown that CNNs with recurrent connections exhibit superior performance when recognising occluded and non-occluded objects ([Spoerer et al., 2017]) and that recurrent connections can help to describe cortical dynamics in early visual cortex ([Kietzmann et al., 2019]). Our plan for the future is to advance upon these important findings by training recurrent connections to predict occluded portions of images as well as behavioural sketches.

Along the same lines, a recently developed network has accurately predicted visual saliency using a similar encoder-decoder network ([Kroner et al., 2020]). Since we know the human visual system’s coding of saliency must be robust to occlusion and clutter, it would be interesting to compare network objectives of predicting occluded visual features and predicting saliency. Such comparisons would allow us to understand whether aspects of these tasks could be shared by the same neuronal pathways.

## Conclusions

We investigated the representational similarity between a specialised artificial network built to reconstruct occluded images and fMRI data obtained while human subjects viewed the same stimuli. Results suggest that low- and mid-level features are present in early visual cortex (V1 and V2). Optimising models to characterise feedback signals in human cortex will improve our understanding about the computations in early visual cortex where we know there is rich top-down information predicting feedforward input, that is currently not captured in feedforward networks. This work points to new experiments in which we challenge AI to replicate as closely as possible brain processing while performing cognitive tasks, testing models explaining memory, visual imagery, and auditory responses in early visual cortex.

## Author Contributions

All authors conceptualised the study. M.S. trained the network. A.T.M. collected and analysed fMRI data. M.S. and A.T.M. computed the RSA analysis. M.S. and L.S.P. wrote the manuscript, and all authors edited the manuscript.

## Footnotes

2 Data will be openly available under EBRAINS knowledge graph after publication.

3 Leaky Relu: *y* = *f* (*x*) = 0.6 ∗ *x* + 0.4 ∗ |*x*| (see figure in Supplementary)

4 The trained model will become openly available under EBRAINS knowledge graph after publication.

5 In this context, a channel - or feature map - is the resulting operation of a single convolution (input image with a single kernel). The number of channels of a layer activation is usually the third dimension after height and width (with 2D input).

