## Supplementary Material for "A Self-Supervised Deep Neural Network for Image Completion Resembles Early Visual Cortex fMRI Activity Patterns for Occluded Scenes"

**Michele Svanera***, **Andrew T. Morgan, Lucy S. Petro, Lars Muckli**
Centre for Cognitive Neuroimaging
Institute of Neuroscience and Psychology
University of Glasgow (UK)
`{name.surname}@glasgow.ac.uk`

December 13, 2020

#### VGG16 representation similarity

Similarity results between VGG16 layer activations and brain data are reported in Figure 1.

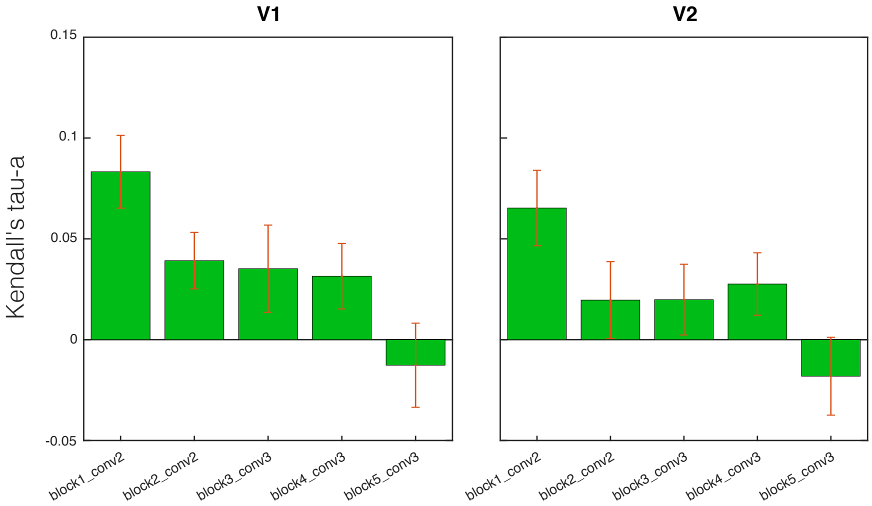


Figure 1: Comparison between brain and VGG16 RDMs. Averaged results across quadrants are shown.

In Fig. 1, results across areas are showing, accordingly with [1, 2], a decreasing of similarity going deeper in the network. The first layer, implementing mainly edge and color contrast detectors (low-level features), is overall the more similar with the brain. From the second, performance is different for every quadrant, which brings a correlation across the entire image to be low. Note that only the first three layers have receptive fields that are separable; from the fourth layer, it is not possible to disentangle quadrants anymore.

*Corresponding author: Michele.Svanera at glasgow.ac.uk

#### Encoder/decoder activation functions

Encoder layers implement rectified linear unit (ReLu) function $y = f(x) = 0.5 * x + 0.5 * |x|$ and decoder leaky ReLu $y = f(x) = 0.6 * x + 0.4 * |x|$. In Figure 2 are shown the two activations.

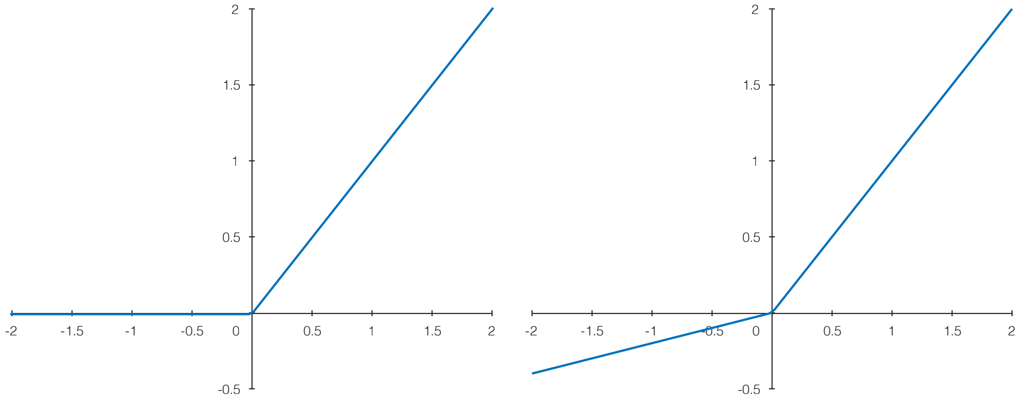


(a) ReLu - Encoder (b) Leaky ReLu - Decoder

Figure 2: Activation functions for (a) encoder and (b) decoder.

#### Training and testing approach

The followed procedure to train our model, takes advantages of two networks: a `generator` (our encoder/decoder in Figure 2), which takes an occluded image as input and produces a reconstructed image in output, and a `discriminator`, which has to detect if the synthetised image is real or fake. In testing, only the generator is used, which, starting from occluded images never seen before (not in the training set), synthesises their fully reconstructed versions. Outlines of training and testing procedures are shown in the Figure 3.

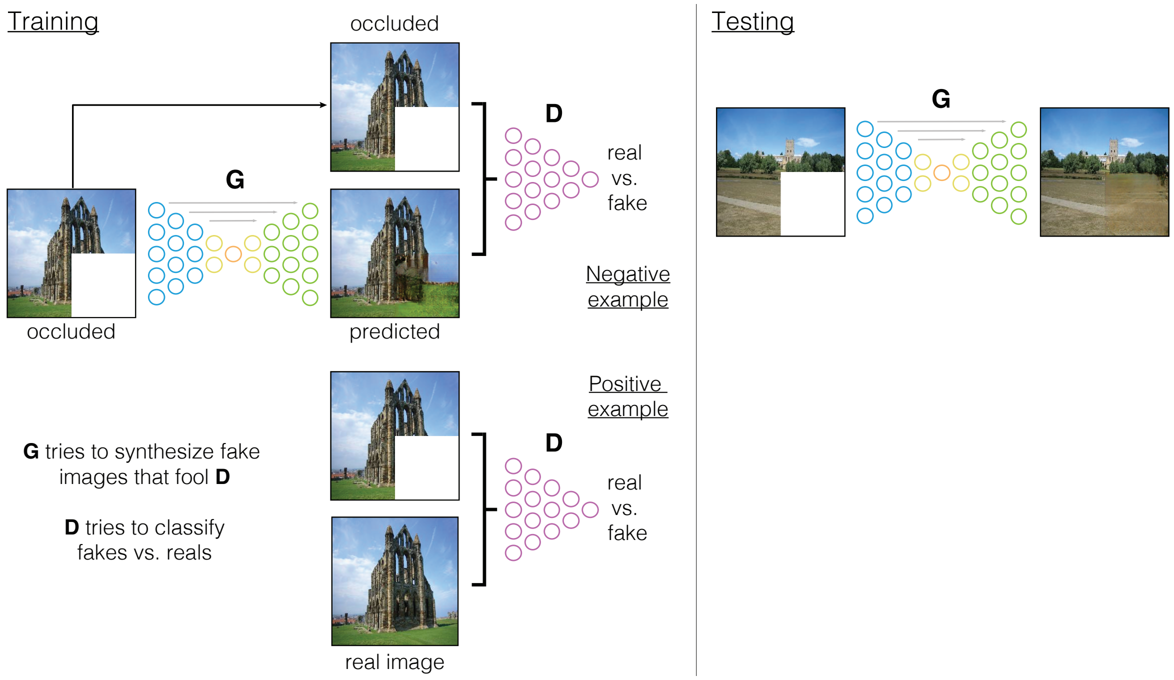


Figure 3: Training and testing procedures of the model. Deliberately inspired by Fig.2 of [3].

#### Experiment images

The 24 images used in the experiment are reported in Figure 4.

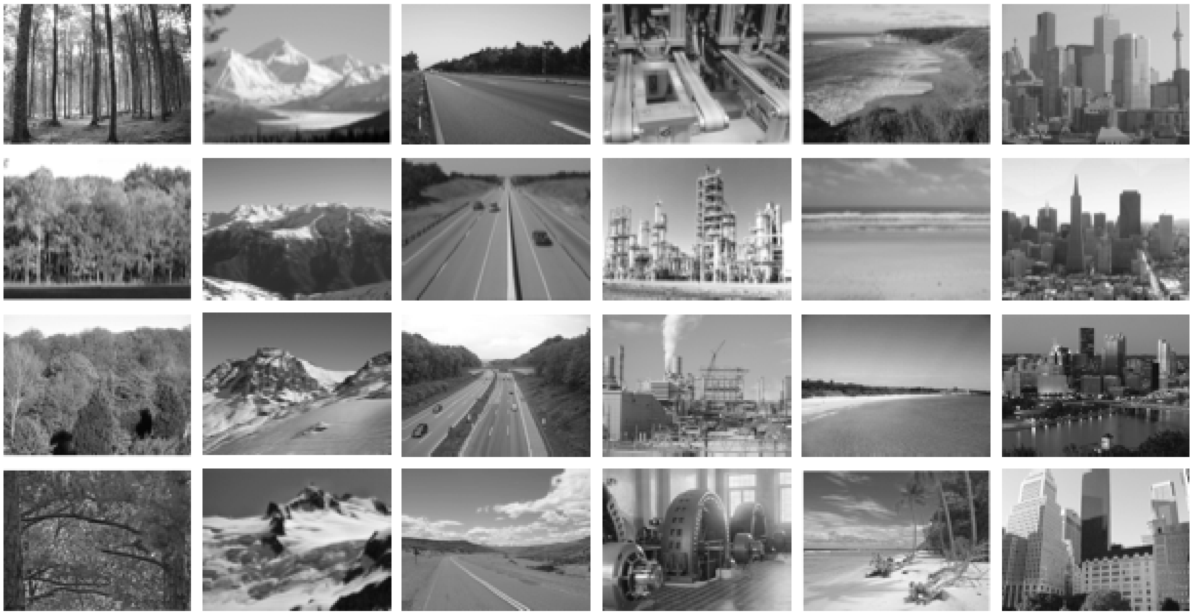

Figure 4: The 24 images (of 6 categories: forests, mountains, highways, industry, beaches, and buildings) used in the experiment. Images are taken from SUN database [4].

#### Population Receptive Field Estimation

In Figure 5, we show the estimation of the Population Receptive Field (pRF), for V1 and V2, across different subjects.

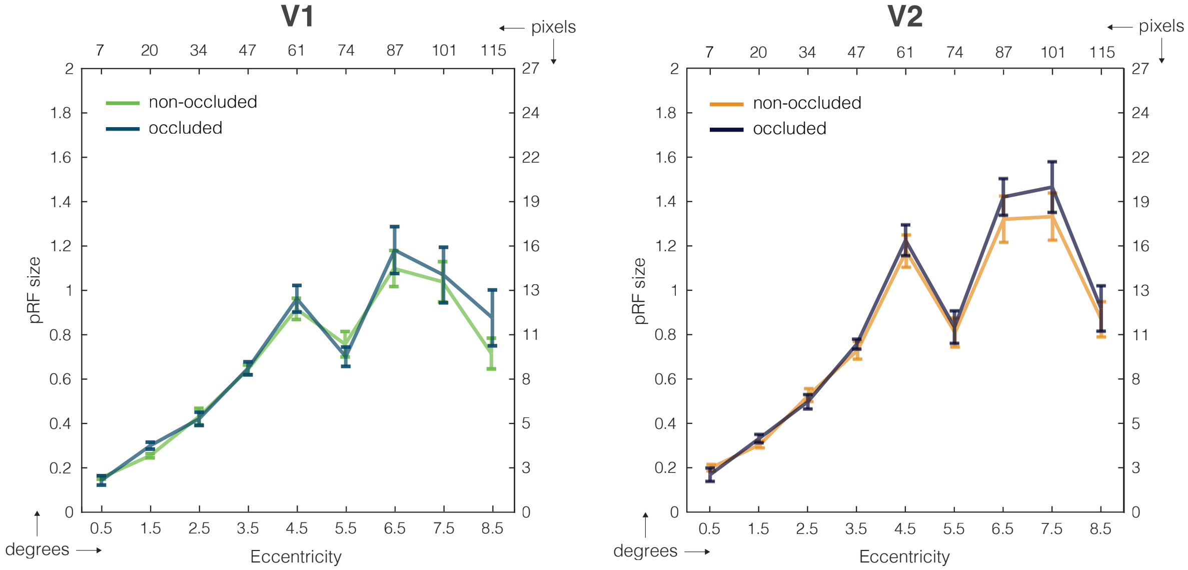


Figure 5: pRF estimation [5]. The first plot is V1, the second is V2. The figures show the pRF size in relation to the distance of the centre of fixation (in degrees of eccentricity and pixels).

Table 1: VGG16 activation dimensions. Total params: $\sim$ 138M

| **Layer (type)** | **Output Shape** | **Receptive field** |
|---|---|---|
| input image | (224, 224, 3) | n.a. |
| block1_conv1 (Conv2D) | (224, 224, 64) | (3, 3) |
| block1_conv2 (Conv2D) | (224, 224, 64) | (5, 5) |
| block1_pool (MaxPooling2D) | (112, 112, 64) | (6, 6) |
| block2_conv1 (Conv2D) | (112, 112, 128) | (10, 10) |
| block2_conv2 (Conv2D) | (112, 112, 128) | (14, 14) |
| block2_pool (MaxPooling2D) | (56, 56, 128) | (16, 16) |
| block3_conv1 (Conv2D) | (56, 56, 256) | (24, 24) |
| block3_conv2 (Conv2D) | (56, 56, 256) | (32, 32) |
| block3_conv3 (Conv2D) | (56, 56, 256) | (40, 40) |
| block3_pool (MaxPooling2D) | (28, 28, 256) | (44, 44) |
| block4_conv1 (Conv2D) | (28, 28, 512) | (60, 60) |
| block4_conv2 (Conv2D) | (28, 28, 512) | (76, 76) |
| block4_conv3 (Conv2D) | (28, 28, 512) | (92, 92) |
| block4_pool (MaxPooling2D) | (14, 14, 512) | (100, 100) |
| block5_conv1 (Conv2D) | (14, 14, 512) | (132, 132) |
| block5_conv2 (Conv2D) | (14, 14, 512) | (164, 164) |
| block5_conv3 (Conv2D) | (14, 14, 512) | (196, 196) |
| block5_pool (MaxPooling2D) | (7, 7, 512) | (212, 212) |
| flatten (Flatten) | (25088) | n.a. |
| fc1 (Dense) | (4096) | n.a. |
| fc2 (Dense) | (4096) | n.a. |
| predictions (Dense) | (1000) | n.a. |

### Model summaries

#### VGG16

In Table 1, VGG16 network details, with activation and receptive field dimensions, are presented. In blue the analysed layers.

#### Encoder/decoder

In Table 2, encoder/decoder network details, with activation and receptive field dimensions, are presented.

### Layer visualisation

#### Layer visualisation encoder/decoder

In Figures 6 to 13 are shown layer activations for `encoder_1` to `encoder_8`. Every Figure corresponds to a specific layer and it has two subplots. **(A)** Once a specific channel of the layer analysed is selected (randomly, the number is reported at the top), the top five activations are shown in column. **(B)** depicts the images where these patches are taken from, with red bounding boxes that indicate the location of the patches in the image.

Table 2: Encoder/decoder activation dimensions. Total params: $\sim$ 228M

| Layer (type) | Output Shape | Receptive field |
|---|---|---|
| input image | (256, 256, 3) | n.a. |
| encoder_1 | (128, 128, 128) | (4, 4) |
| encoder_2 | (64, 64, 256) | (10, 10) |
| encoder_3 | (32, 32, 512) | (22, 22) |
| encoder_4 | (16, 16, 1024) | (46, 46) |
| encoder_5 | (8, 8, 1024) | (94, 94) |
| encoder_6 | (4, 4, 1024) | (190, 190) |
| encoder_7 | (2, 2, 1024) | n.a. |
| encoder_8 | (1, 1, 1024) | n.a. |
| decoder_8 | (2, 2, 1024) | n.a. |
| decoder_7 | (4, 4, 1024) | n.a. |
| decoder_6 | (8, 8, 1024) | n.a. |
| decoder_5 | (16, 16, 1024) | n.a. |
| decoder_4 | (32, 32, 512) | n.a. |
| decoder_3 | (64, 64, 256) | n.a. |
| decoder_2 | (128, 128, 128) | n.a. |
| output image | (256, 256, 3) | n.a. |

##### Layer visualisation VGG16

In Figures 14 to 18 are shown layer activations for `block1_conv2`, `block2_conv2`, `block3_conv3`, `block4_conv3`, and `block5_conv3`. Every Figure corresponds to a specific layer and it has two subplots. **(A)** Once a specific channel of the layer analysed is selected (randomly, the number is reported at the top), the top five activations are shown in column. **(B)** depicts the images where these patches are taken from, with red bounding boxes that indicate the location of the patches in the image.

[3] Phillip Isola, Jun-Yan Zhu, Tinghui Zhou, and Alexei A Efros. Image-to-image translation with conditional adversarial networks. *CVPR*, 2017.

[4] Jianxiong Xiao, James Hays, Krista A Ehinger, Aude Oliva, and Antonio Torralba. Sun database: Large-scale scene recognition from abbey to zoo. In *Computer vision and pattern recognition (CVPR), 2010 IEEE conference on*, pages 3485–3492. IEEE, 2010.

[5] Serge O. Dumoulin and Brian A. Wandell. Population receptive field estimates in human visual cortex. *NeuroImage*, 39(2):647 – 660, 2008.

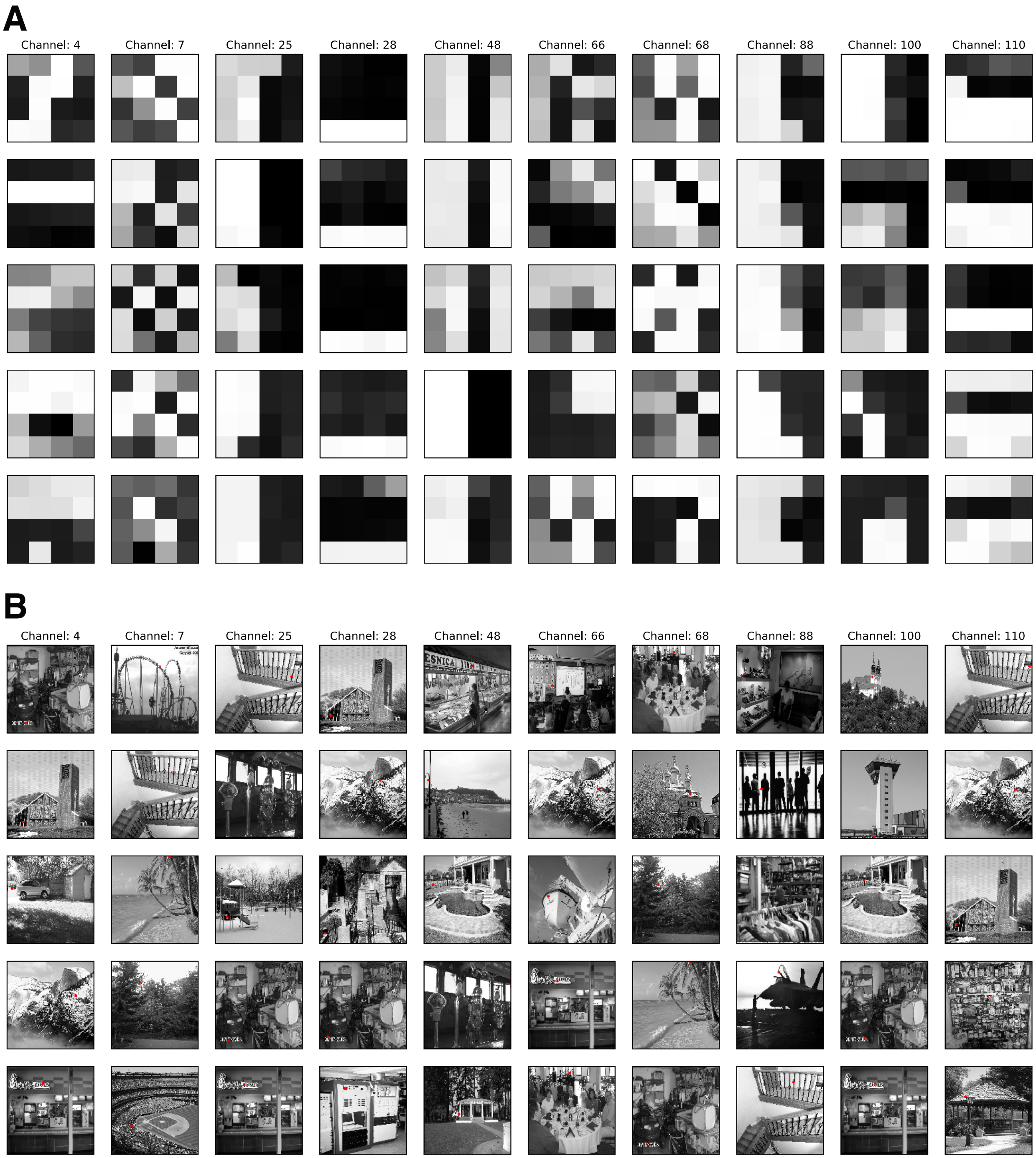


Figure 6: Layer `encoder_1` max activations. Every column is a channel. **(A)** Receptive fields, in the image space, that provide the five largest activations for a specific channel. **(B)** Original images are provided.

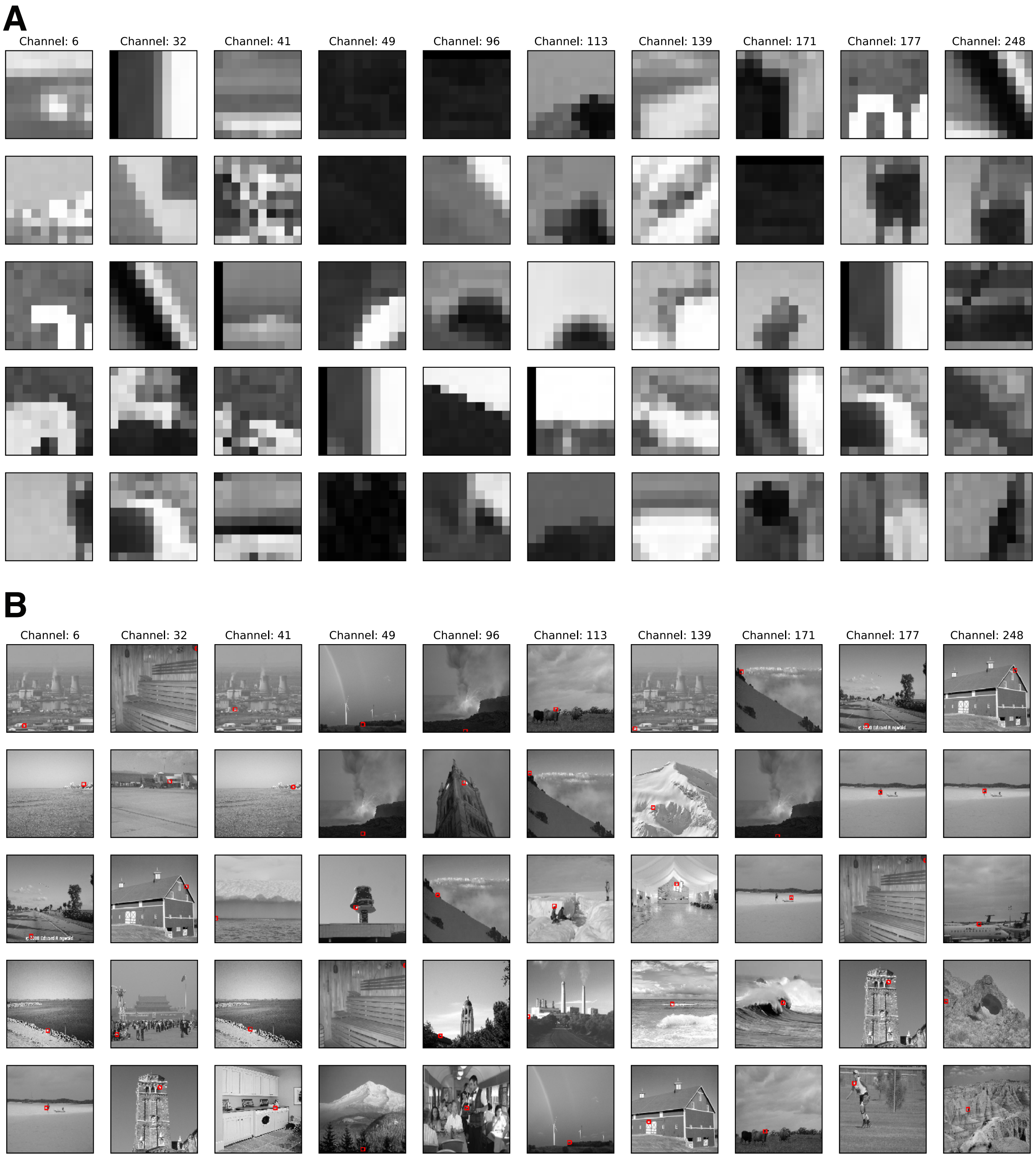


Figure 7: Layer `encoder_2` max activations. Every column is a channel. **(A)** Receptive fields, in the image space, that provide the five largest activations for a specific channel. **(B)** Original images are provided.

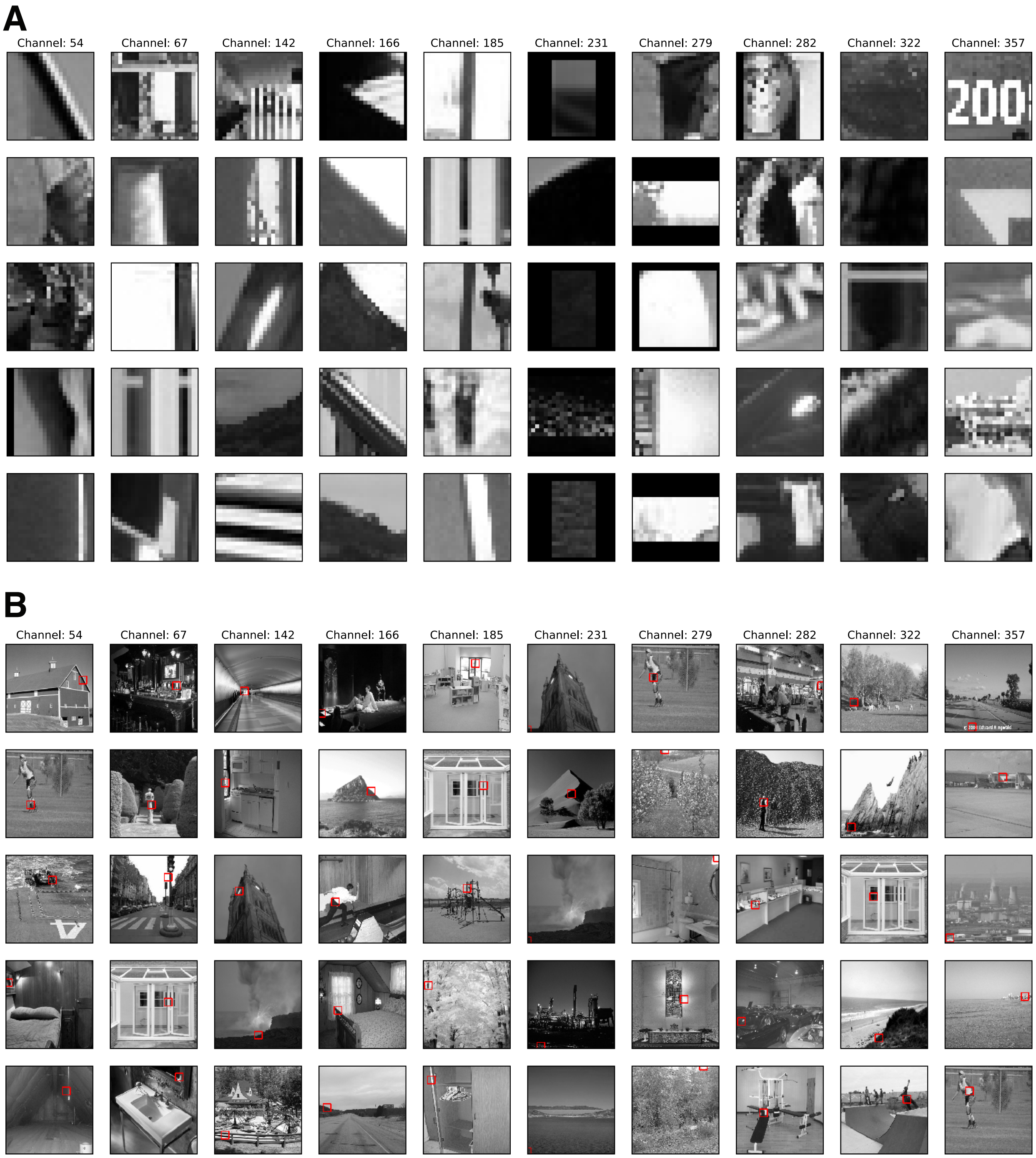


Figure 8: Layer `encoder_3` max activations. Every column is a channel. **(A)** Receptive fields, in the image space, that provide the five largest activations for a specific channel. **(B)** Original images are provided.

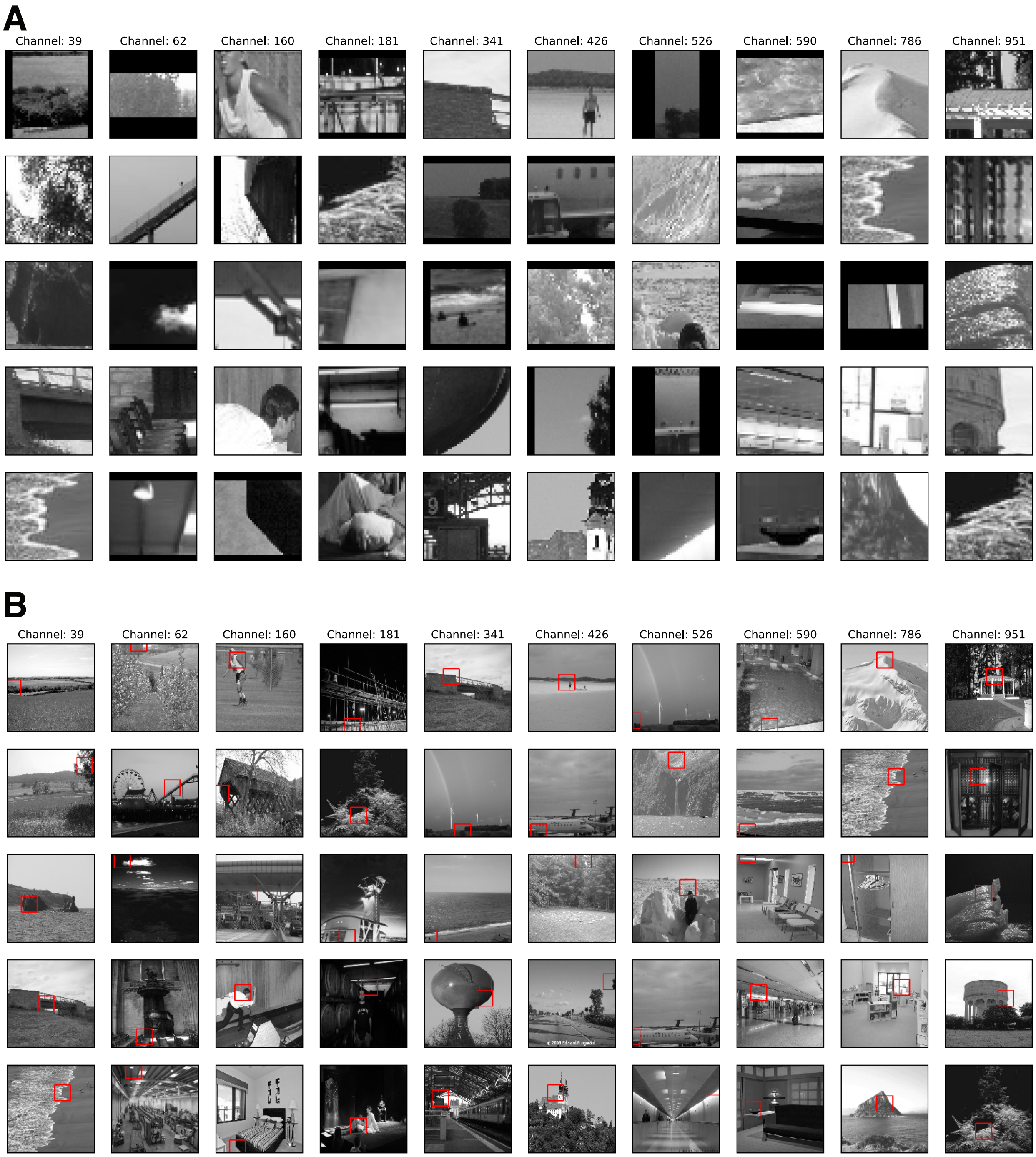


Figure 9: Layer `encoder_4` max activations. Every column is a channel. **(A)** Receptive fields, in the image space, that provide the five largest activations for a specific channel. **(B)** Original images are provided.

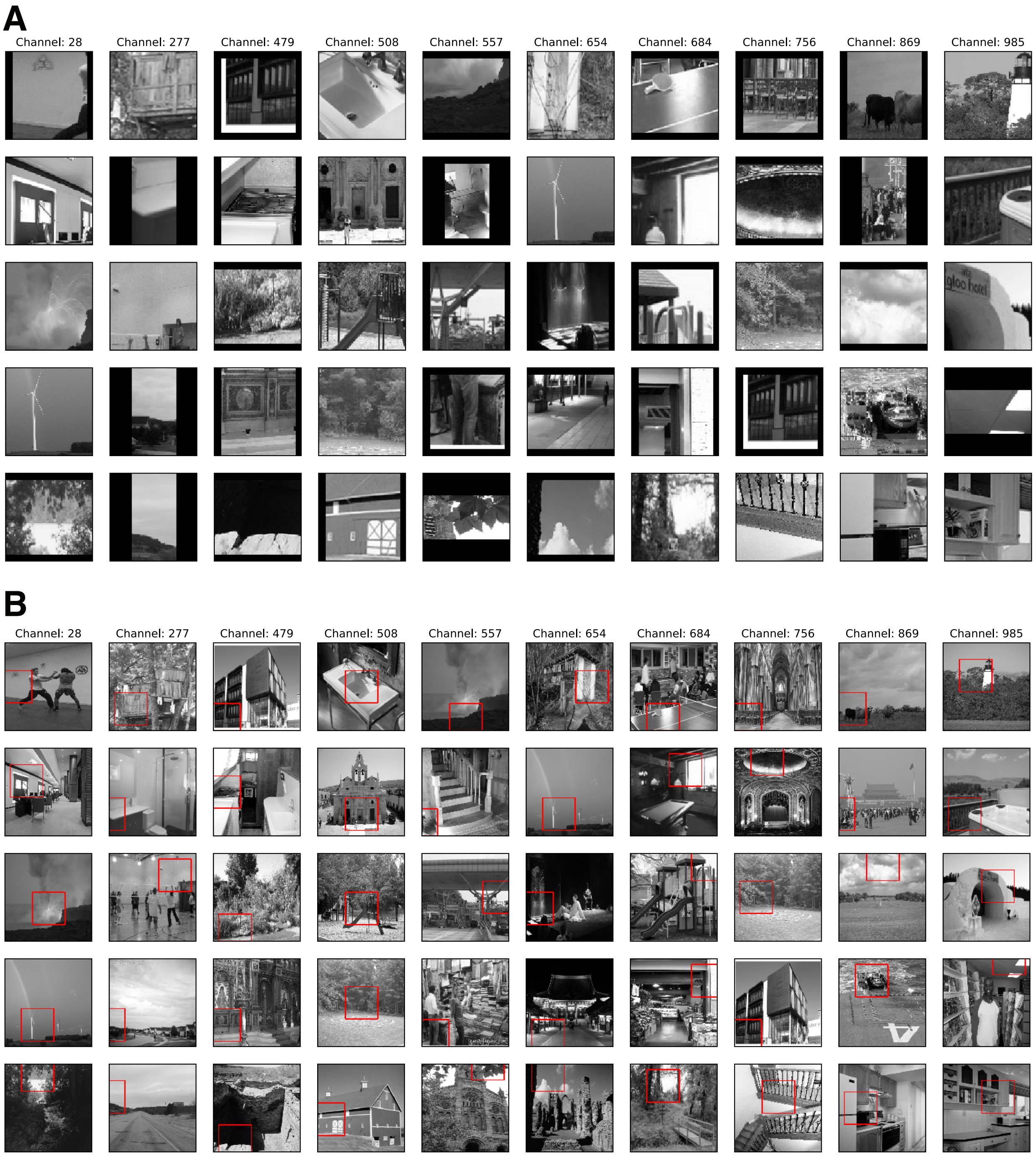


Figure 10: Layer `encoder_5` max activations. Every column is a channel. **(A)** Receptive fields, in the image space, that provide the five largest activations for a specific channel. **(B)** Original images are provided.

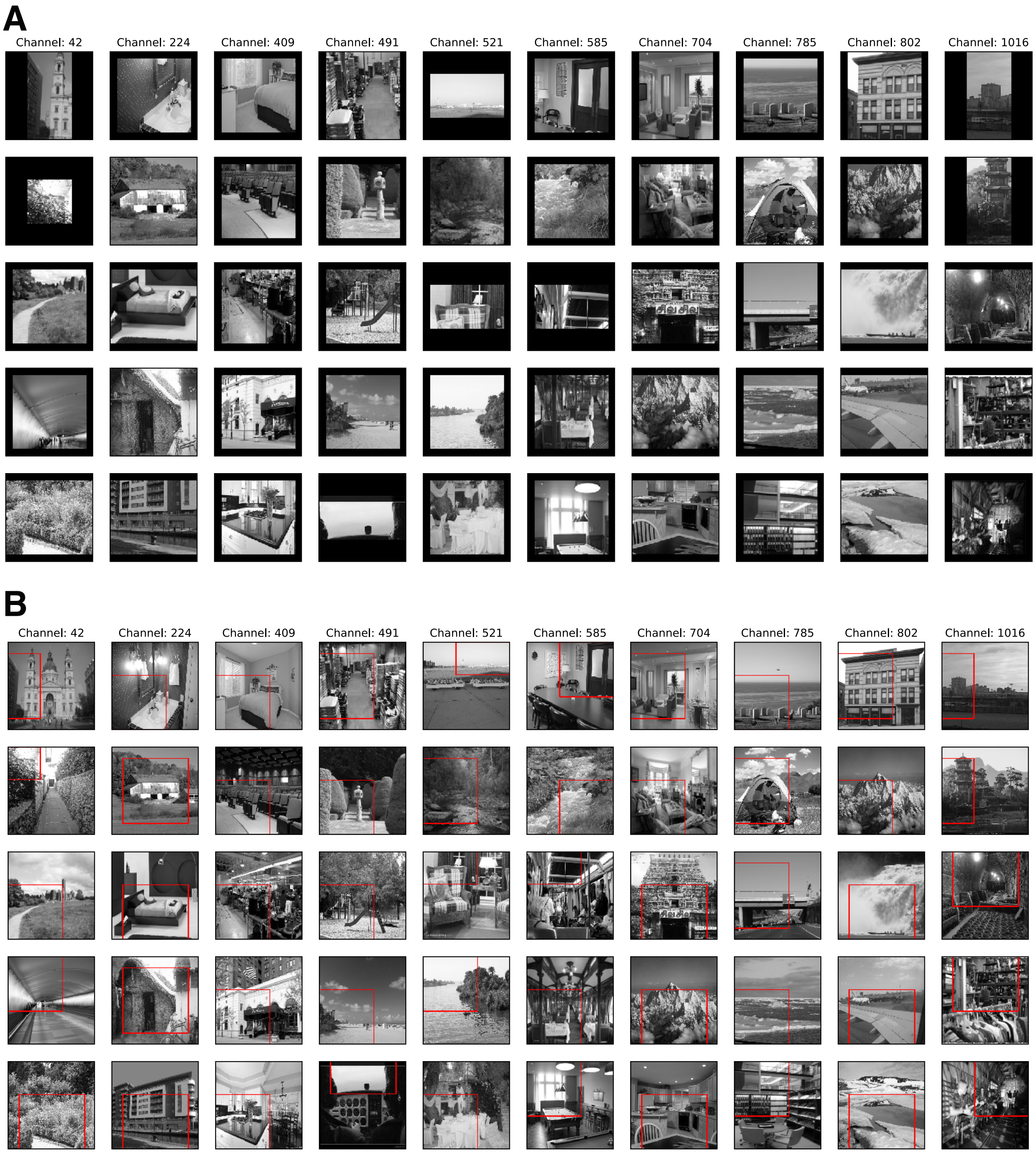


Figure 11: Layer `encoder_6` max activations. Every column is a channel. **(A)** Receptive fields, in the image space, that provide the five largest activations for a specific channel. **(B)** Original images are provided.

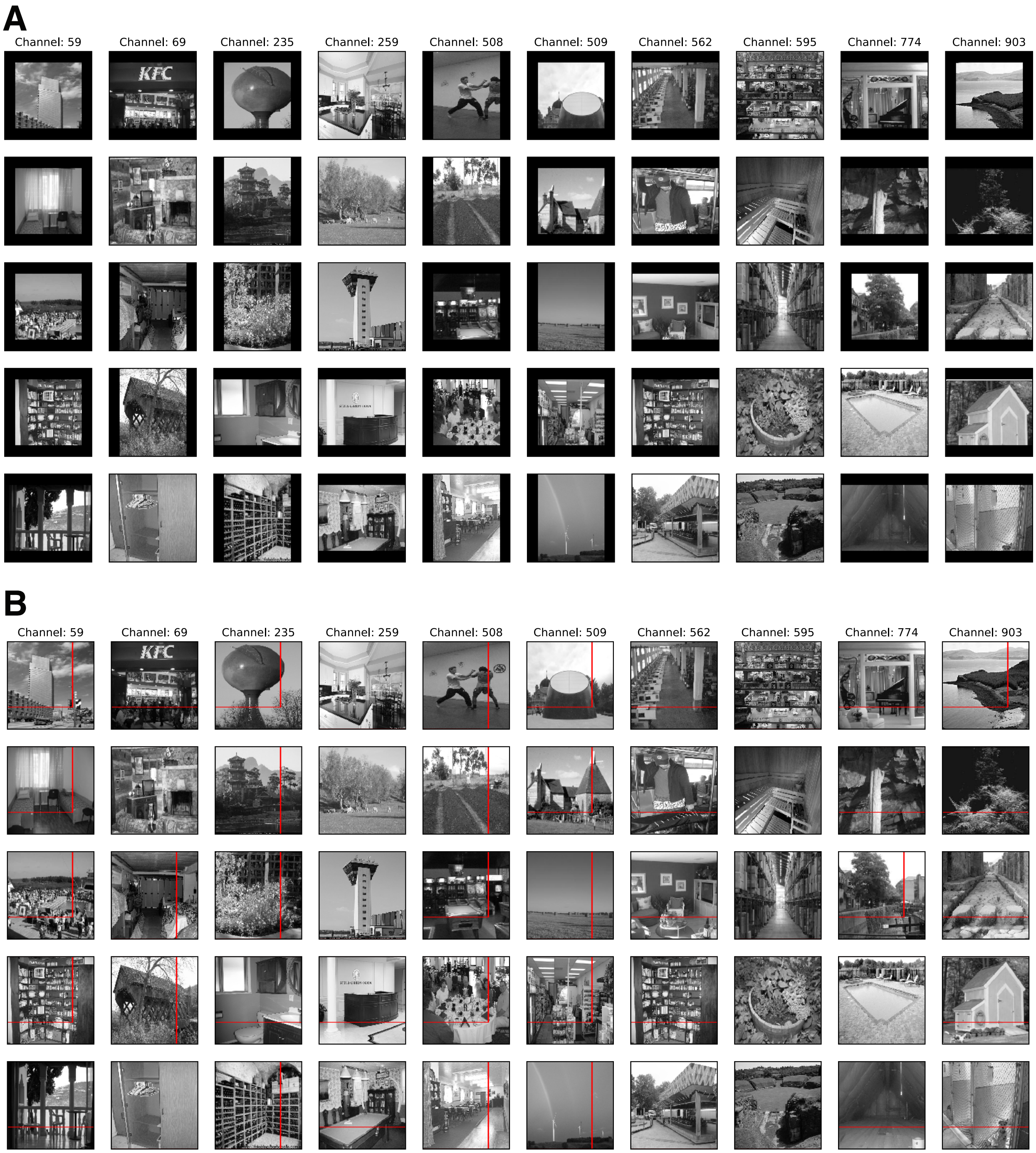


Figure 12: Layer `encoder_7` max activations. Every column is a channel. **(A)** Receptive fields, in the image space, that provide the five largest activations for a specific channel. **(B)** Original images are provided.

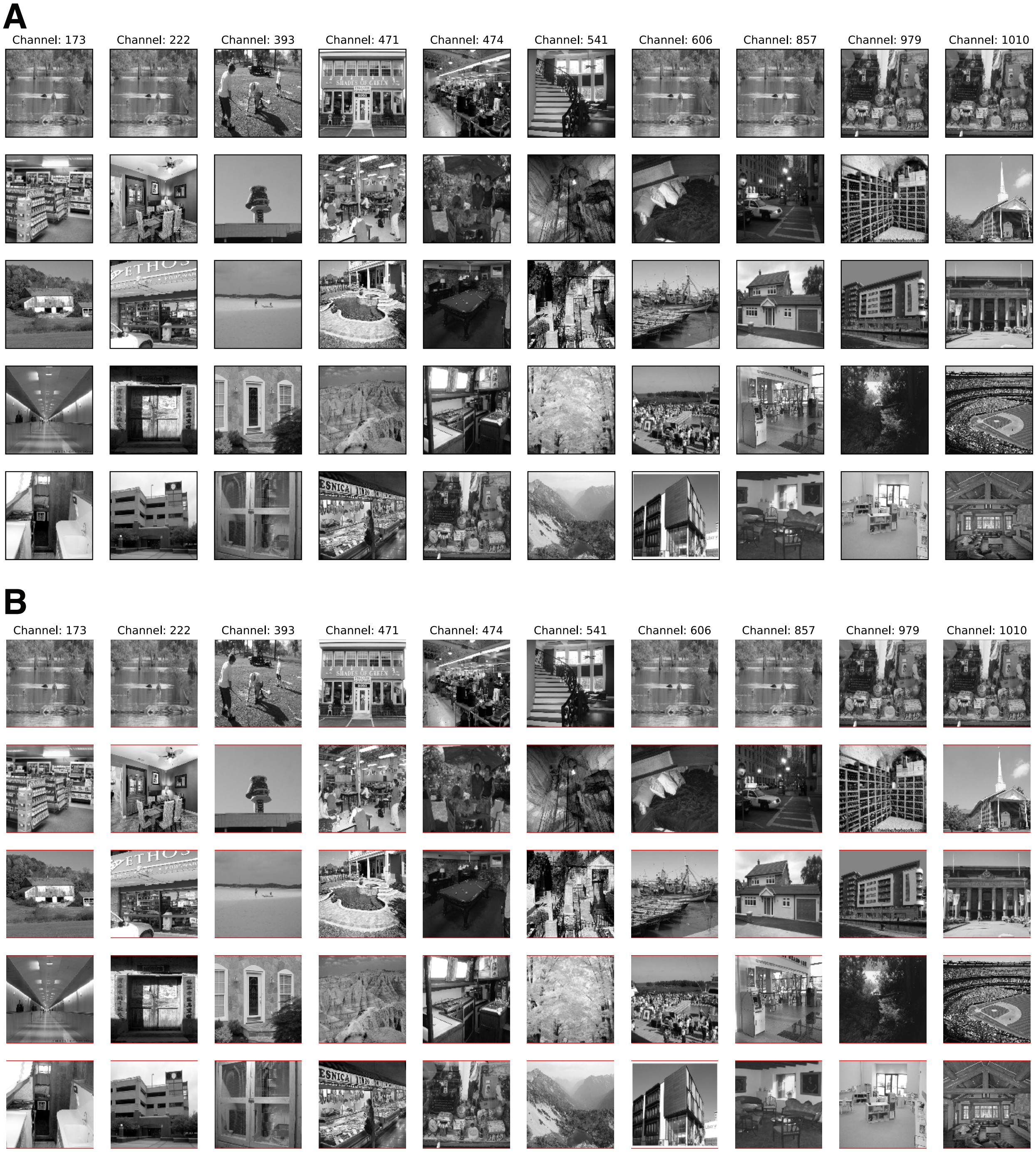


Figure 13: Layer `encoder_8` max activations. Every column is a channel. **(A)** Receptive fields, in the image space, that provide the five largest activations for a specific channel. **(B)** Original images are provided.

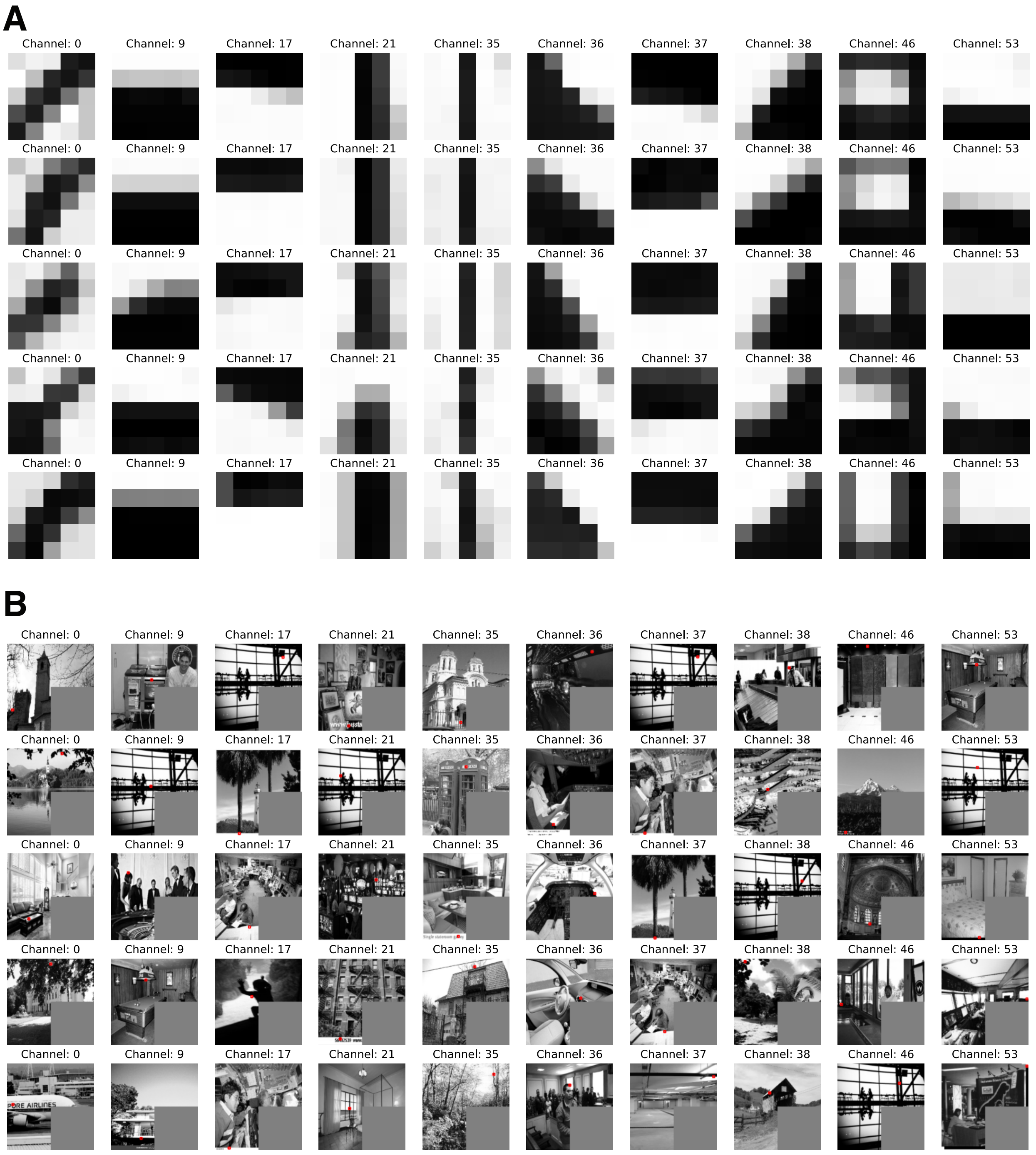


Figure 14: Layer `block1_conv2` max activations. Every column is a channel. **(A)** Receptive fields, in the image space, that provide the five largest activations for a specific channel. **(B)** Original images are provided.

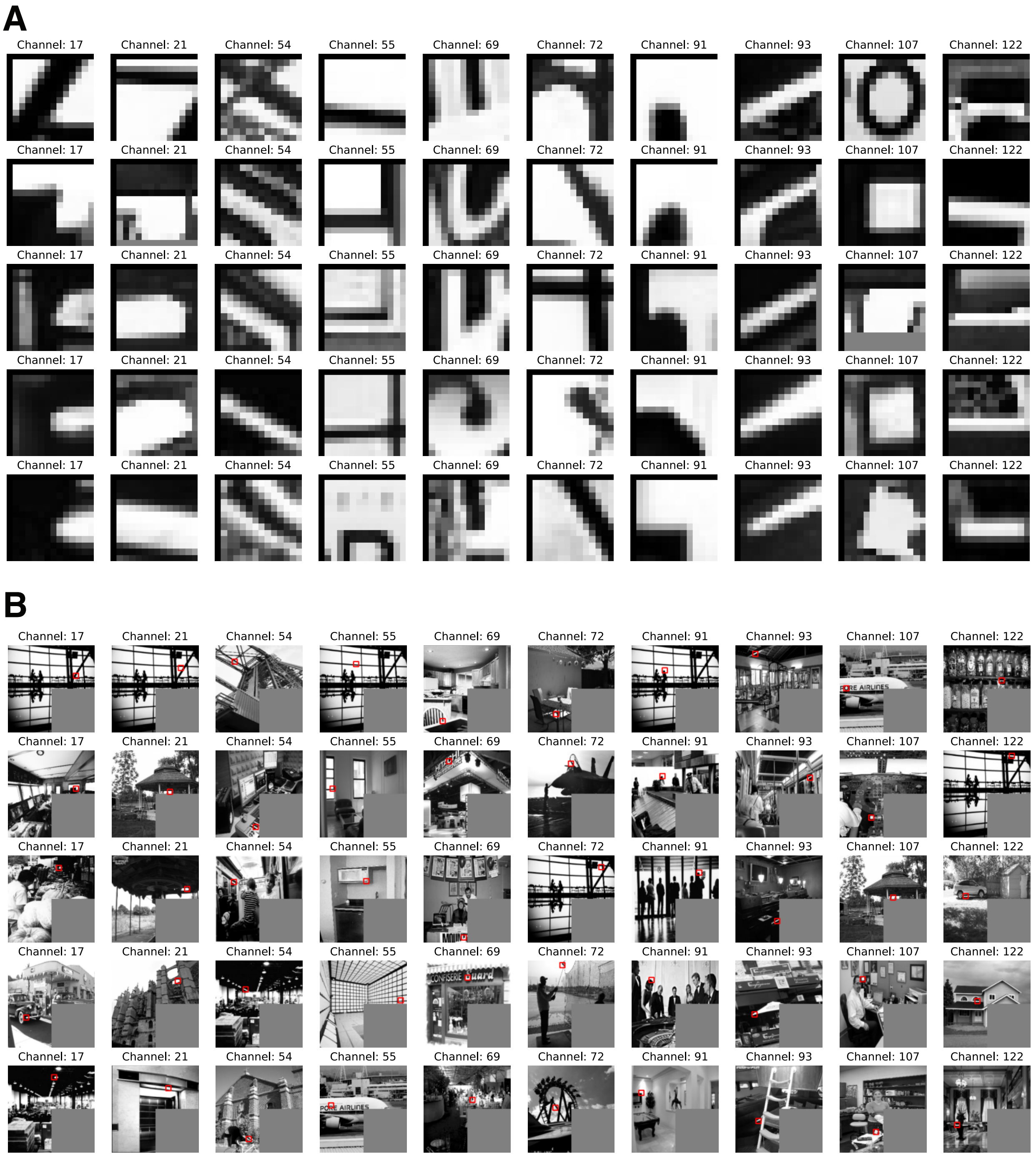


Figure 15: Layer `block2_conv2` max activations. Every column is a channel. **(A)** Receptive fields, in the image space, that provide the five largest activations for a specific channel. **(B)** Original images are provided.

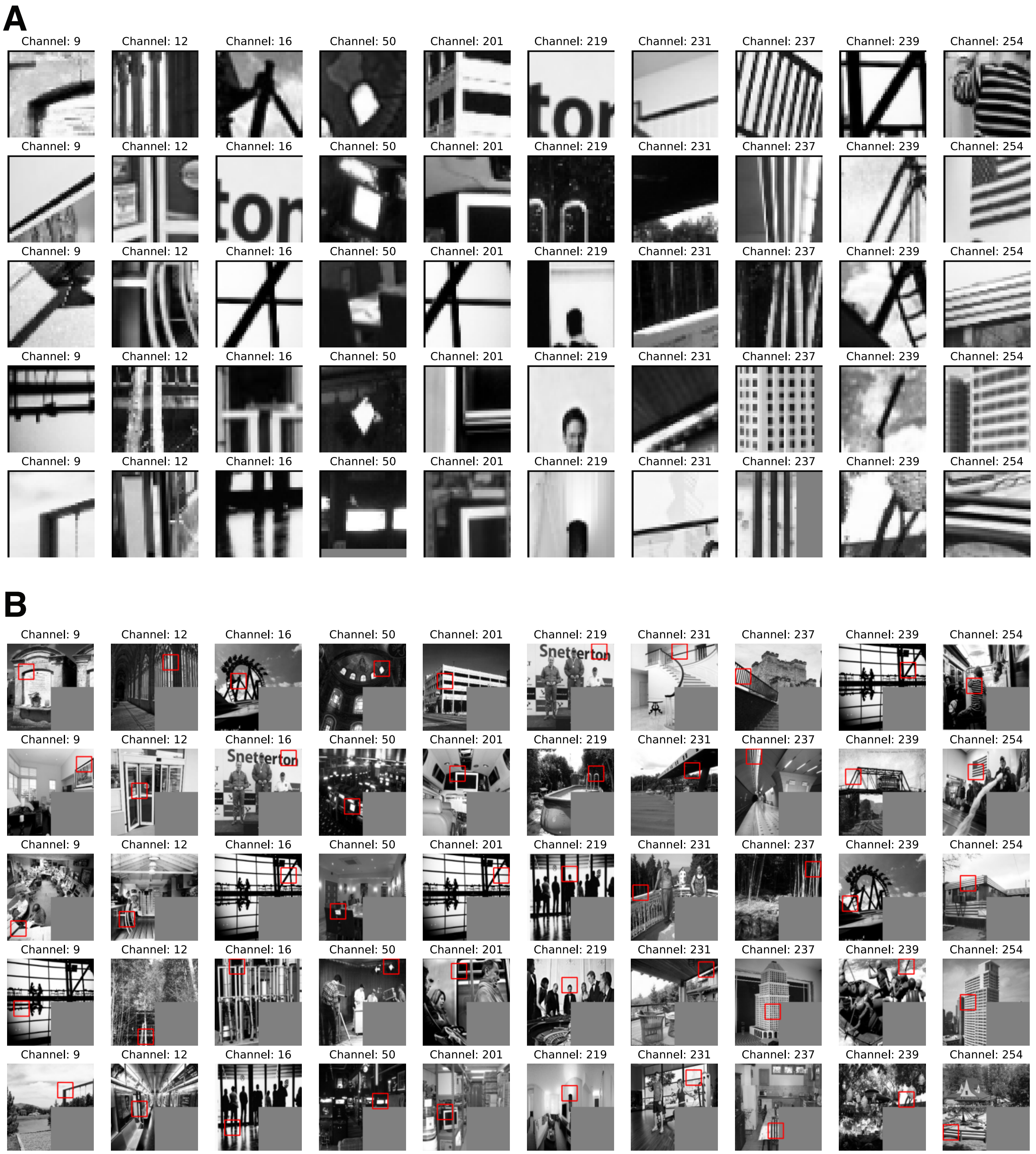


Figure 16: Layer `block3_conv3` max activations. Every column is a channel. **(A)** Receptive fields, in the image space, that provide the five largest activations for a specific channel. **(B)** Original images are provided.

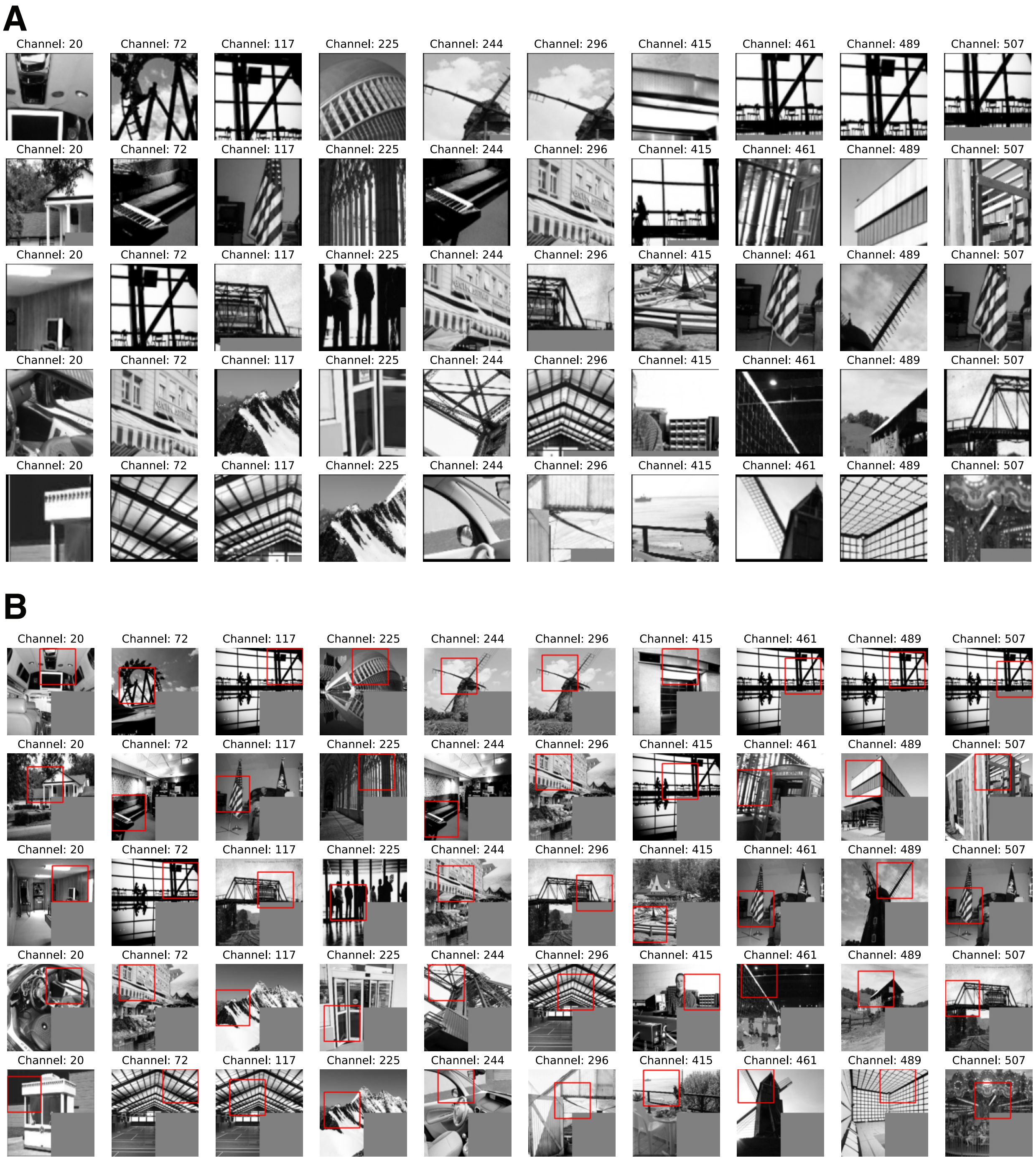


Figure 17: Layer `block4_conv3` max activations. Every column is a channel. **(A)** Receptive fields, in the image space, that provide the five largest activations for a specific channel. **(B)** Original images are provided.

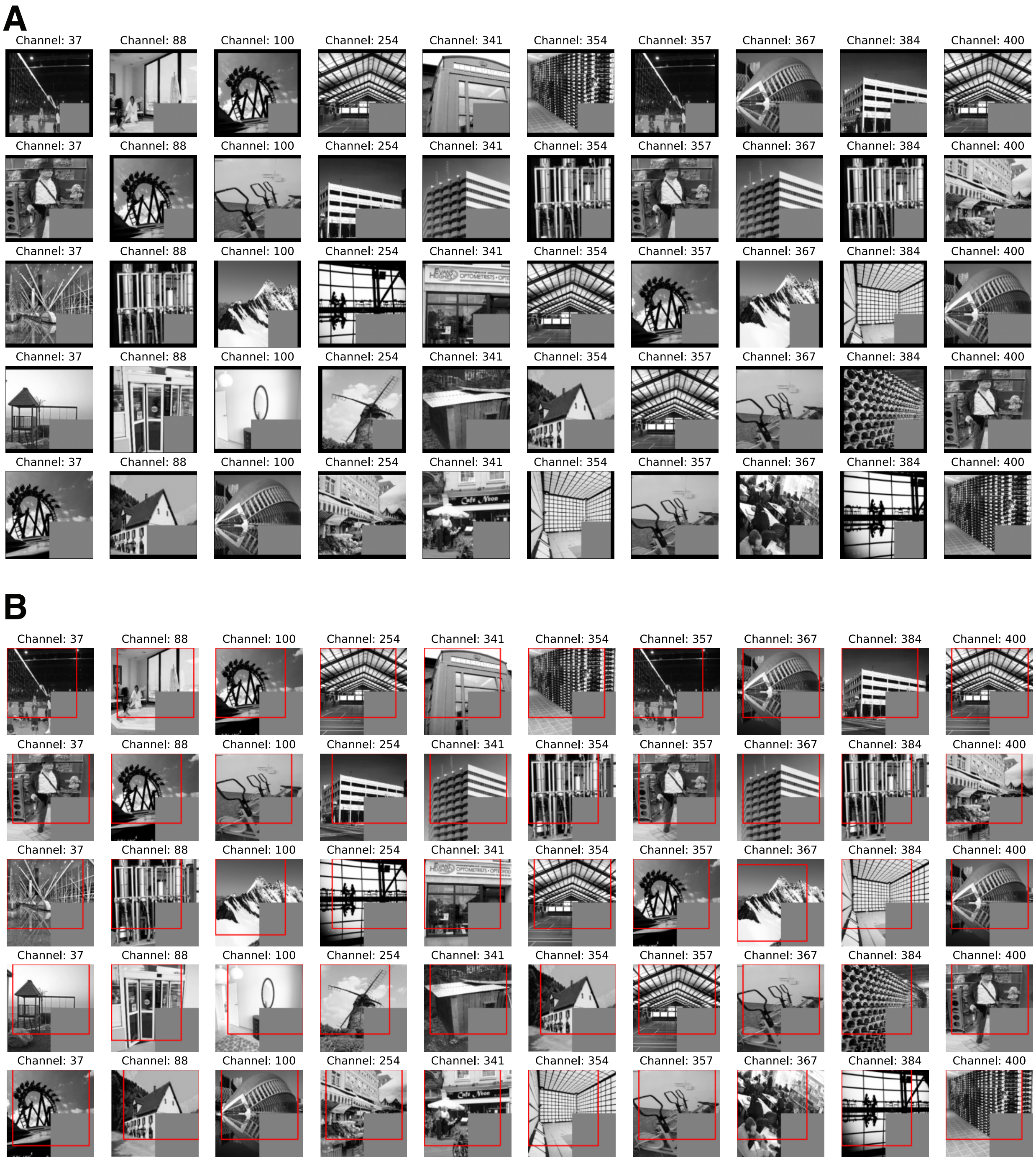


Figure 18: Layer `block5_conv3` max activations. Every column is a channel. **(A)** Receptive fields, in the image space, that provide the five largest activations for a specific channel. **(B)** Original images are provided.
